# Regulation of cell–nanoparticle interactions through mechanobiology

**DOI:** 10.1101/2024.07.07.599665

**Authors:** Marco Cassani, Francesco Niro, Helena Durikova, Sofia Morazzo, Daniel Pereira-Sousa, Soraia Fernandes, Jan Vrbsky, Jorge Oliver-De La Cruz, Simon Klimovic, Jan Pribyl, Tomas Loja, Petr Skladal, Frank Caruso, Giancarlo Forte

## Abstract

Bio–nano interactions have been extensively explored in nanomedicine to develop selective delivery strategies, reduce systemic toxicity, and minimize therapeutic dosing requirements. To enhance the delivery of nanocarriers to cancer cells and improve the therapeutic efficiency and clinical translation of nanomedicines, numerous nanomaterials with diverse and tunable properties have been developed. However, the limited clinical translation of nanoparticle-based therapies, largely due to issues associated with poor targeting and therapeutic delivery, requires a deeper understanding of the biological phenomena underlying cell–nanoparticle interactions. In this context, herein we investigate the molecular and cellular mechanobiology parameters that control such interactions. We demonstrate that the pharmacological inhibition or the genetic ablation of the key mechanosensitive component of the Hippo pathway, i.e., yes-associated protein, enhances nanoparticle internalization by 1.5-fold. Importantly, this phenomenon occurs independently of nanoparticle properties, such as size, or cell properties such as surface area, substrate adhesion, and stiffness. Our study reveals that the internalization of nanoparticles in target cells can be controlled by modulating cell mechanosensing pathways, potentially ultimately enhancing nanoparticle delivery and nanotherapy specificity.

## Introduction

In recent decades, understanding the interactions between nanomaterials and biological systems has become a primary goal in nanomedicine, aiming to design nanomaterials (including nanoparticles, NPs) that can efficiently engage with living cells.^1–4^ While efforts have mostly focused on engineering the physicochemical properties of nanoparticles, such as size, shape, stiffness, and surface chemistry, with the goal to enhance their interaction with cells and improve drug delivery, the role of intracellular molecular pathways in bio–nano interactions has often been overlooked.^5–7^ For instance, it is known that the stiffness of nanomaterials can influence their association with cell membranes, leading to different outcomes depending on their size and composition.^8–10^ Similarly, surface charge influences nanoparticle–cell membrane interactions, with positively charged nanoparticles being internalized more readily than negatively charged ones.^11, 12^ Despite the significant progress achieved over the past two decades in understanding bio–nano interactions, there is a need to further elucidate mechanisms governing nanoparticle interactions with biological environments.^13, 14^ Unveiling the processes responsible for nanomaterial–cell interactions at the molecular level may shed new light on nanoparticle transport within specific cells.^15^ Consequently, a paradigm shift in which intracellular molecular processes redefine the interpretation of bio–nano interactions may lead to novel strategies for improving nanoparticle delivery and overall applications in nanomedicine. In this context, cell mechanics has emerged as a promising area of investigation, revealing its role in regulating cell–nanoparticle interactions.^16, 17^ The mechanical state of cells has been shown to influence the degree of nanoparticle binding and internalization, paving the way to the *mechanotargeting* of primary or metastatic cancer cells through nanoparticles.^18^ More recently, the reduction of tumor stiffness obtained through the inhibition of focal adhesion kinase and concomitant reduction in the deposition of extracellular matrix (ECM) components was found to promote the delivery of lipid nanoparticles and transfection efficiency in cancer cells.^19^

Recently, mechanotherapeutics has emerged as a new class of drugs and treatments targeting mechanically activated pathways involved in various pathologies.^20^ Targeting mechanosensing pathways has resulted in promising outcomes in guiding cell fate and modulating cell function.^21^ The components of such pathways are responsible for controlling the expression of genes related to cell migration and survival, and cancer malignant progression through the recruitment of specific transcription factors.^22, 23^ Among the key players in cell mechanosensing, yes-associated protein (YAP) has emerged as a central regulator of mechanotransduction in cancer cells.^24^ YAP, a mechanoactivated protein acting as the downstream effector of the Hippo pathway, is frequently dysregulated in cancer and contributes to cell proliferation, migration, survival, and immune evasion.^25–27^ We reported that YAP deletion in cancer cells leads to significant changes in cell shape and morphology, substrate adhesion, and the perception of mechanical cues generated within the surrounding microenvironment.^28^ More recently, our work revealed that YAP regulates cell–nanoparticle interactions and the delivery of nanodrugs by influencing cellular mechanical properties, the expression of genes involved in endocytic pathways, and the deposition of ECM components.^29^ Specifically, we demonstrated that the genetic or pharmacological inhibition of YAP significantly enhances nanoparticle internalization in the triple-negative breast cancer (TNBC) cell line CAL51, which is characterized by high YAP expression and transcriptional activity.^28^ To deepen our understanding of this phenomenon, in the present study, we investigated the role of YAP on nanoparticle uptake in a different cell model, i.e., HEK 293T, which displays a lower YAP expression and transcriptional activity than CAL51.^29^ Despite HEK 293T cells showing lower dependence on YAP activity in terms of mechanical properties such as adhesion, shape, and stiffness, our findings further elucidate the role of YAP in cell–nanoparticle interactions. Additionally, our research highlights the potential for reducing YAP expression to enhance nanoparticle uptake. Thus, by modulating the activity of YAP, we sought to determine the role of the Hippo effector and its associated mechanoregulated pathways in directing the outcome of nanoparticle association with cells.

## Result and discussion

### YAP depletion has minimal effects on the mechanical and physical properties of HEK 293T cells

Recently, we demonstrated that the knockdown of YAP in TNBC cell line CAL51 significantly influenced cell morphology, adhesion, and stiffness.^29^ YAP depletion resulted in an overall reduction in cell area, substrate adhesion, and membrane rigidity, which was accompanied by a reduction in ECM deposition and an increase in the uptake of nanoparticles and nanodrugs. Herein, to understand the role of mechanobiology effectors on nanoparticle association with cells, we investigate the role of YAP in cells characterized by a lower protein expression, thus rendering them less dependent on YAP activity (Fig. S1).^29^ By minimizing the influence of other factors, such as membrane stiffness and cell surface area, which may significantly influence cell–nanoparticle interactions, we sought to determine the unique role of YAP in this process.

Using CRISPR/Cas9 technology, we generated a stable YAP-deficient mutant HEK 293T cell line^29^ and confirmed YAP depletion through western blot analysis (Fig. 1a). At the transcriptional level, reverse transcription-quantitative polymerase chain reaction (RT-qPCR) showed a marked reduction in the mRNA levels of YAP accompanied by a decrease in the expression of connective tissue growth factor (CTGF), one of the main transcriptional targets of YAP (Fig. 1b). The physical and mechanical properties of the HEK 293T cells remained unaltered after YAP depletion, as determined by atomic force microscopy (AFM) (Fig. 1c). Likewise, cell surface area, as assessed by membrane extension and actin coverage, was unaffected (Fig. 1d,e). Furthermore, confocal images revealed that wild-type (WT) HEK and YAP −/− HEK share the same morphology and distribution of the focal adhesion protein vinculin (Fig. 1f.; Fig. S2).

**Fig. 1.**
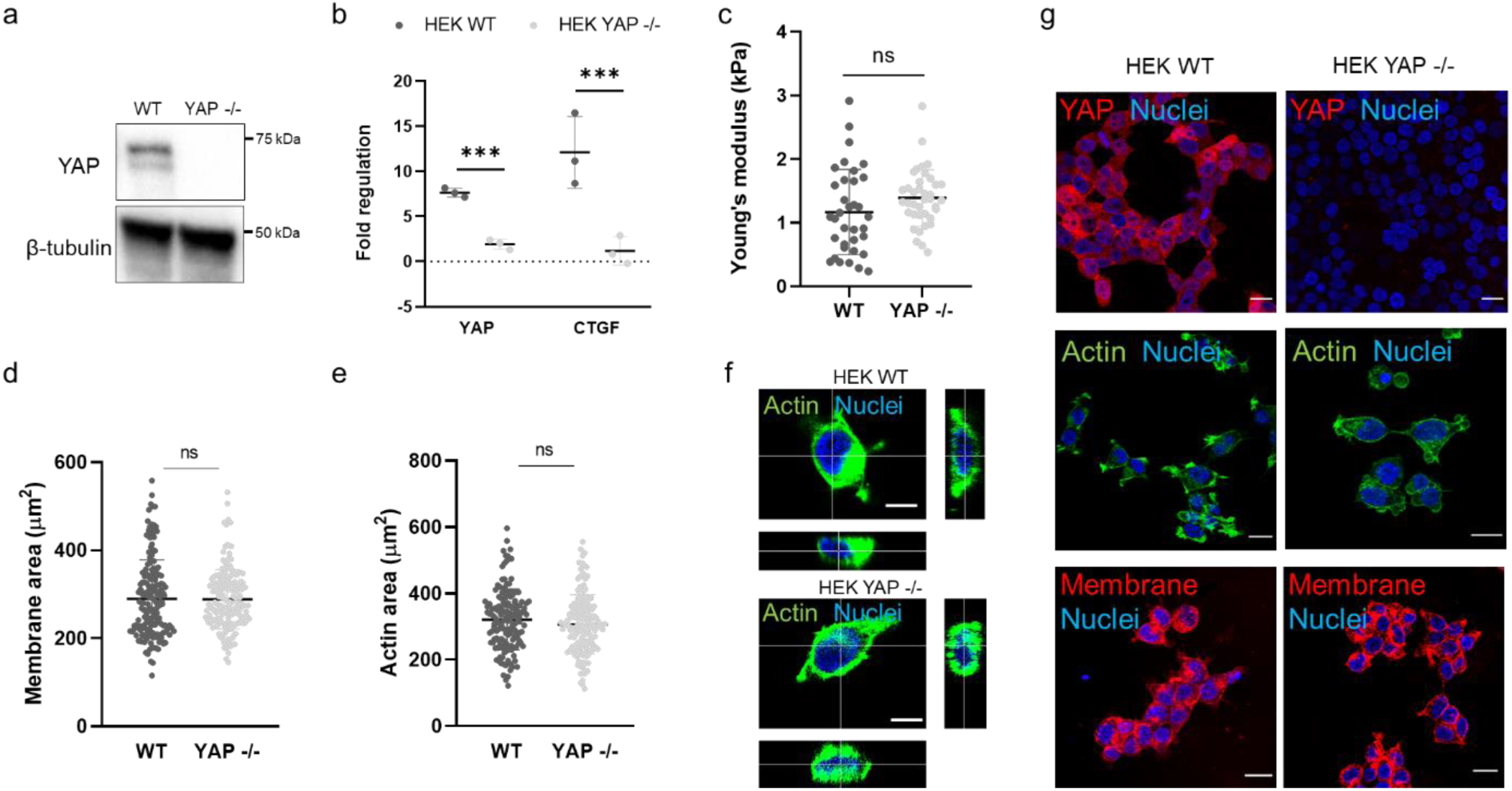
YAP depletion does not influence the adhesion, mechanics, or morphology properties of HEK cells. (a) Western blot analysis showing the levels of YAP protein in WT and YAP −/− HEK cells. β-Tubulin was used for protein loading normalization. (b) RT-qPCR analysis of YAP and CTGF in WT and YAP −/− HEK cells. Statistical analysis was performed using multiple *t*-test; *n* = 3; \*\*\**p* < 0.001. (c) Dot plots of the Young’s moduli of WT and YAP −/− HEK cells, as measured by AFM. Statistical analysis was performed using unpaired *t*-test with Welch’s correction; ns, nonsignificant. (d) Dot plot analysis of the total membrane area of WT and YAP −/− HEK cells. Cells were stained with Alexa Fluor 488-labeled wheat germ agglutinin (WGA-488); *n* > 100. Statistical analysis was performed using unpaired *t*-test with Welch’s correction; ns, nonsignificant. (e) Dot plot analysis of the surface area of WT and YAP −/− HEK cells, as calculated on the basis of the total actin coverage of the cells. Cells were stained with Alexa Fluor 488-labeled phalloidin (Pha-488, green); *n* > 100. Statistical analysis was performed using unpaired *t*-test with Welch’s correction; ns, nonsignificant. (f) Three-dimensional (3D) reconstruction of WT and YAP −/− HEK cells. Cells were stained with 4′,6′-diamidino-2-phenylindole (DAPI) and Pha-488. Scale bars: 10 μm. (g) Representative confocal images depicting YAP expression in WT and YAP −/− HEK cells. Cells were stained for YAP (Alexa Fluor 555, red, top), actin (Pha-488, green, middle), membrane (wheat germ agglutinin–Alexa Fluor 647 conjugate (WGA-647), red, bottom) and nuclei were counterstained with DAPI. Scale bars: 20 μm.

These results indicate that YAP depletion in HEK 293T cells influences the expression of target genes and the YAP-related transcriptional activity of the cells but has minimal effects on the physical and mechanical properties of the cells.

### YAP activity controls the binding and internalization of nanoparticles in HEK 293T cells

As shown in our previous work,^29^ YAP activity hampered the internalization of nanoparticles in TNBC cell line CAL51; this phenomenon was also observed in our preliminary studies in HEK 293T cells. However, while the regulation of this process in CAL51 could be attributed to the significant effects that YAP displayed on cell morphology, adhesion, membrane structure and ECM deposition, this may differ in HEK 293T as these effects were only marginal (see Fig. 1).

To examine how this process is regulated in HEK cells, carboxylated polystyrene (PS) spherical nanoparticles of 200 and 900 nm in diameter (denoted as PS200 and PS900, respectively) that were labeled with carboxytetramethylrhodamine were used to treat WT and YAP −/− HEK cells (incubation period of 4 h), and cell–nanoparticle interactions were investigated using confocal microscopy and flow cytometry. The confocal analysis revealed that the PS nanoparticles preferentially bound to YAP −/− HEK cells than to WT cells (Fig. 2a–c). This effect was independent of the nanoparticle size and was confirmed by flow cytometry, which showed higher nanoparticle uptake ratios in YAP −/− HEK than in WT cells (Fig. 2d). Importantly, no change in membrane stiffness was observed after nanoparticle binding, indicating that the cell– nanoparticle interactions did not influence the mechanical properties of the cells (Fig. 2e,f). Furthermore, *z*-stack confocal images confirmed the more effective internalization of the nanoparticles by YAP −/− cells, with a higher number of particles found to bind and colocalize with the cell membrane in the absence of YAP (Fig. 2g,h). As expected, this phenomenon was also observed at later time-points (Fig. S3). Collectively, our results show increased nanoparticle association and internalization in YAP −/− HEK cells despite the marginal effect that YAP depletion has on the physical and mechanical properties of HEK 293T cells.

**Fig. 2.**
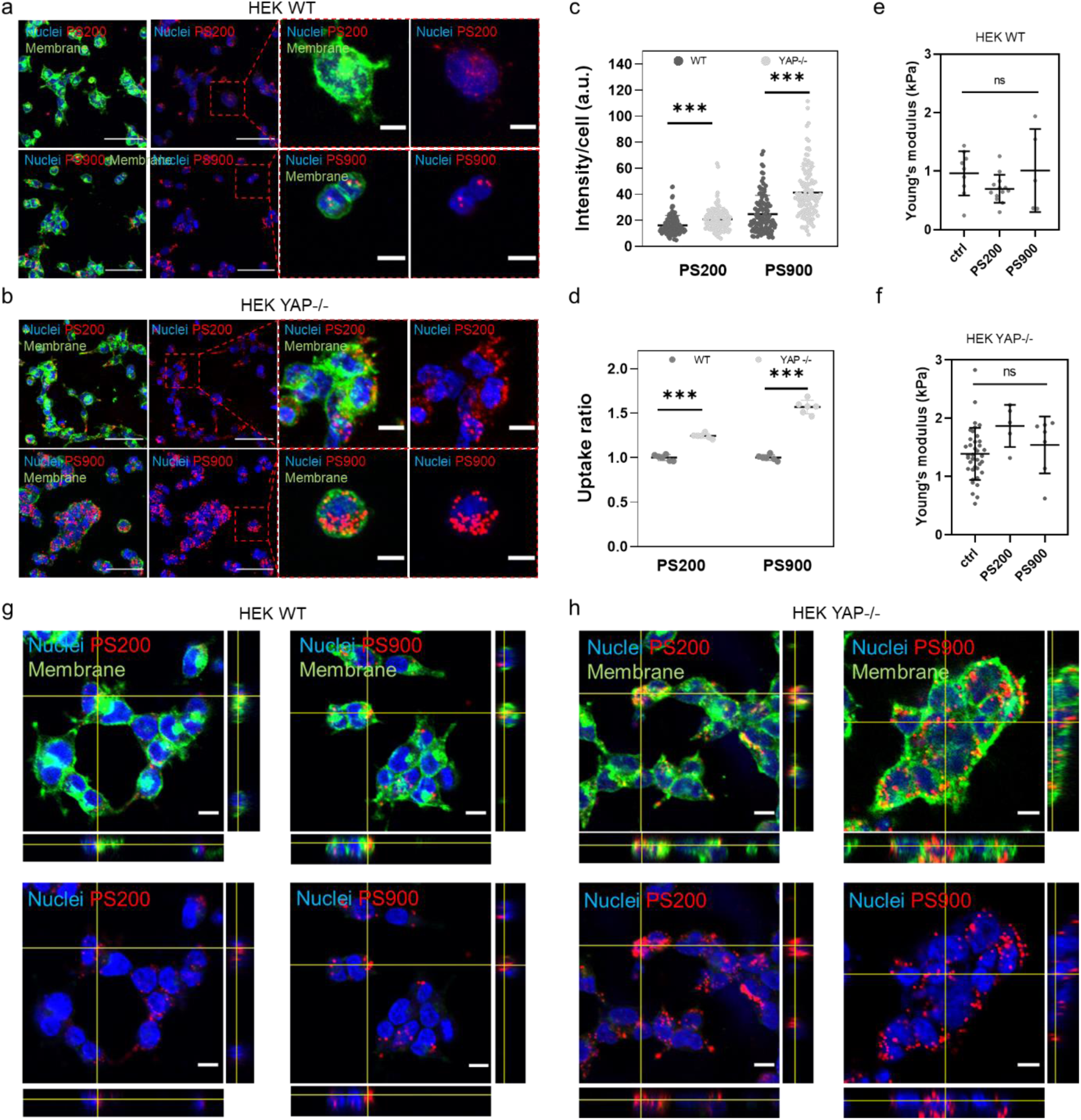
YAP regulates nanoparticle association with HEK 293T cells. (a, b) Representative confocal images of WT (a) and YAP −/− (b) HEK cells after incubation for 4 h with PS200 or PS900. Cells are stained with WGA-488 (green) and/or DAPI (blue). Magnified images of the regions within the red dashed boxes are also shown. Scale bars: 50 and 10 μm for the lower and higher magnification images, respectively. (c) Nanoparticle intensity per cell profile after incubation of PS200 or PS900 with WT and YAP −/− HEK cells for 4 h. Statistical analysis was performed using unpaired *t*-test; *n* > 100; \*\*\**p* < 0.001. (d) Uptake ratios of PS200 and PS900 in WT or YAP −/− HEK cells after incubation for 4 h. Statistical analysis was performed using two-way analysis of variance (ANOVA) followed by Sidak’s multiple comparison test; *n* = 6; \*\*\**p* < 0.001. (e, f) Young’s moduli of WT (e) and YAP −/− (f) HEK cells after incubation for 4 h with PS200 or PS900, as measured by AFM. Statistical analysis was performed using Kruskal–Wallis one-way ANOVA followed by Dunn’s multiple comparisons test; ns, nonsignificant. (g, h) Orthogonal views of *z*-projections of WT (g) and YAP −/− (h) HEK cells after incubation for 4 h with PS200 or PS900. Cells were stained with WGA-488 (green) and DAPI (blue). Scale bars: 10 μm.

### YAP transcriptional activity governs membrane organization in HEK 293T cells

To explore the processes responsible for the different interactions with nanoparticles, we conducted an unbiased RNA sequencing (RNA-seq) analysis and evaluated changes induced by YAP depletion in the HEK 293T transcriptional landscape. The analysis revealed a total of 503 differentially expressed genes in YAP −/− HEK cells, with 297 of the expressed genes being downregulated and 206 of them being upregulated following YAP depletion (Fig. S4). Notably, the most represented gene ontology (GO) annotations were associated with membrane organization, with 53% of genes downregulated in YAP −/− HEK involved in membrane organization (Fig. 3a). Among the genes downregulated in YAP −/− HEK and involved in membrane organization, we identified *CAV1* (log_2_ fold-change (log_2_FC) 1.6), *RhoA* (log_2_FC 1.01), and *TLCD2* (log_2_FC 1.1) (Fig. 3b,c). *CAV1* is a negative regulator of endocytosis, which has been demonstrated to inhibit dynamin-dependent raft-mediated endocytosis, possibly playing a role as endocytosis inhibitor along the caveolae pathway.^30^ *RhoA*, member of the Rho small GTPases family, is responsible for reducing fluid phase uptake and endocytosis in a process that involves membrane organization and is dependent on its activity.^31^ Finally, *TLCD2* is a transmembrane protein involved in the incorporation of polyunsaturated fatty acids into phospholipids, playing a role in regulating membrane fluidity and organization.^32^

**Fig. 3.**
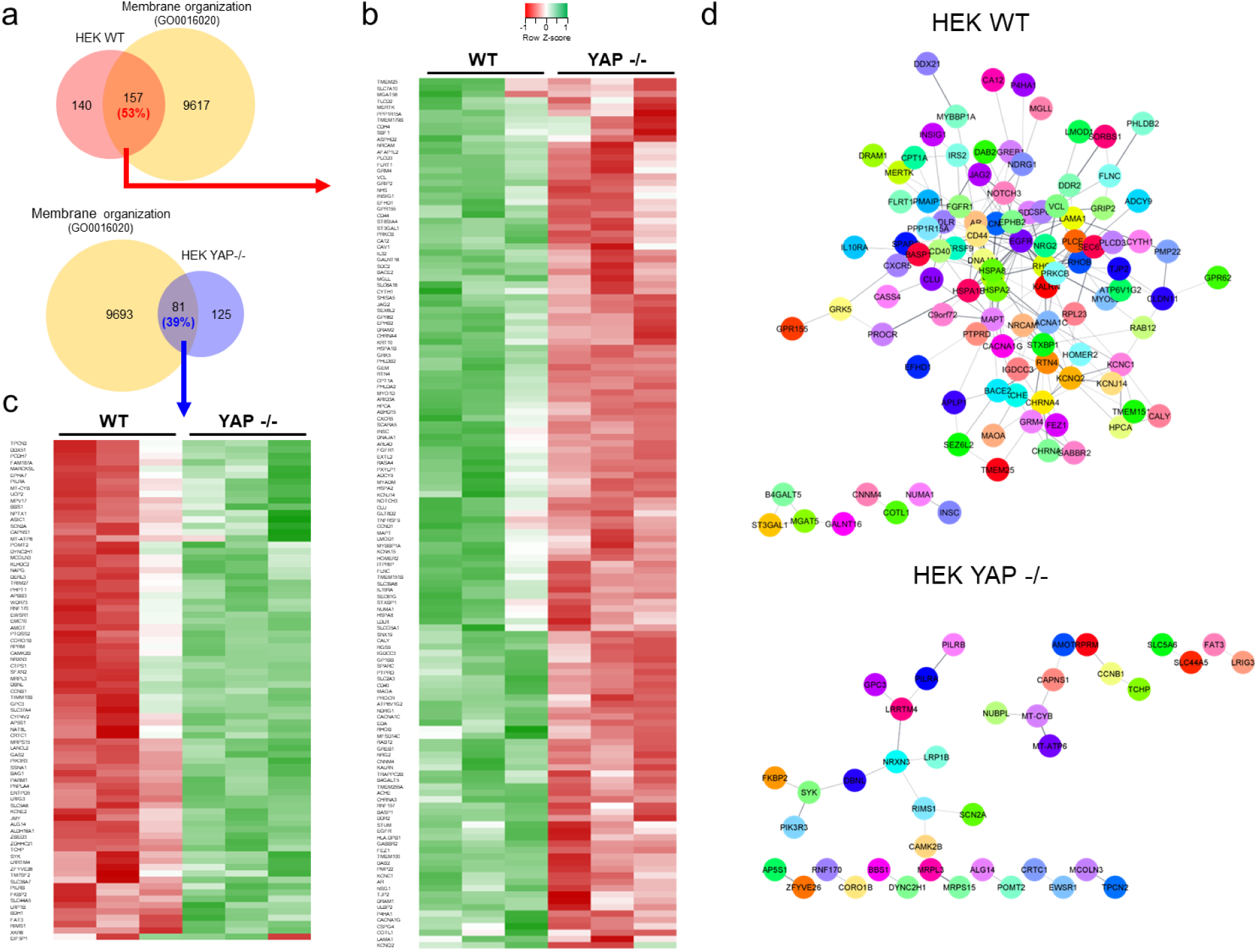
YAP depletion in HEK cells alters the expression of genes related to membrane organization. (a) Venn diagram showing the overlap between genes significantly downregulated in WT and YAP −/− HEK cells and belonging to the membrane organization network (GO0016020), as obtained by RNA-seq. (b, c) Heatmaps of the relative expression of significantly regulated genes associated with the membrane organization network, significantly downregulated (b) or upregulated (c) in YAP −/− HEK cells. *n* = 3 (P adj < 0.05, log2FC > ǀ1ǀ). (d) STRING PPI network of differentially expressed proteins involved in membrane organization in WT and YAP −/− HEK cells obtained from Cytoscape (P adj < 0.05, log2FC > ǀ1ǀ, confidence cutoff 0.4).

In addition, STRING PPI analysis yielded a highly clustered network (cluster coefficient 0.26) containing 156 nodes and 229 edges for HEK WT. In contrast, only 81 nodes and 27 edges were identified in YAP-depleted cells (Fig. 3d; Fig. S5). WT HEK cells showed a higher number of interconnections than YAP −/− HEK cells, indicating that YAP regulates the expression of gene whose protein forms an intricated network in controlling membrane organization.

The cytosolic retention of YAP is commonly linked to protein turnover and degradation pathway alternative to the Hippo pathway, which involves large tumor suppressor homolog 1/2 (LATS1/2) phosphorylation and proteasomal degradation.^33^ However, recent research has unveiled the relationship between YAP, the cell membrane, and the endocytic machinery in the cytoplasm.^34, 35^ These interactions may reveal novel and unexpected functions of the cytoplasmic pool of YAP in direct interaction with membrane proteins, vesicles, and organelles. Despite these possibilities, the precise role of YAP in these processes remains unclear.

Collectively, the present findings suggest that interactions with nanoparticles are promoted in YAP-depleted cells owing to the dysregulation of an interconnected network of key regulators of membrane organization and assembly.

### Overexpression of YAP in HEK 293T cells impairs their ability to bind to nanoparticles

Given that HEK 293T cells displayed low cytoplasmic YAP expression that could not explain the differences in nanoparticle association found in CAL51 cell line,^29^ we conducted a series of experiments involving HEK 293T cells in which we induced increased YAP expression in the WT or knockout (KO) background. Cells were transfected with a plasmid carrying a transcriptionally hyperactive form of YAP, known as YAP S6A. This protein variant contains specific mutations that convert the serine residues S61, S109, S127, S128, S131, S136, S164, and S381 into alanine residues.^36^ The accumulation of YAP in the cell nuclei, and consequently its function, is mostly controlled by a cascade of kinases of the Hippo pathway that phosphorylate YAP on serine residues, promoting its degradation in the cytosol and limiting its cotranscriptional activity (Fig. 4a).^33^ The mutations carried by YAP S6A render the protein nonphosphorylatable and resistant to inactivation and degradation, resulting in the activation and translocation of the protein into the nucleus, where it can function as a transcriptional coactivator (Fig. 4b).

**Fig. 4.**
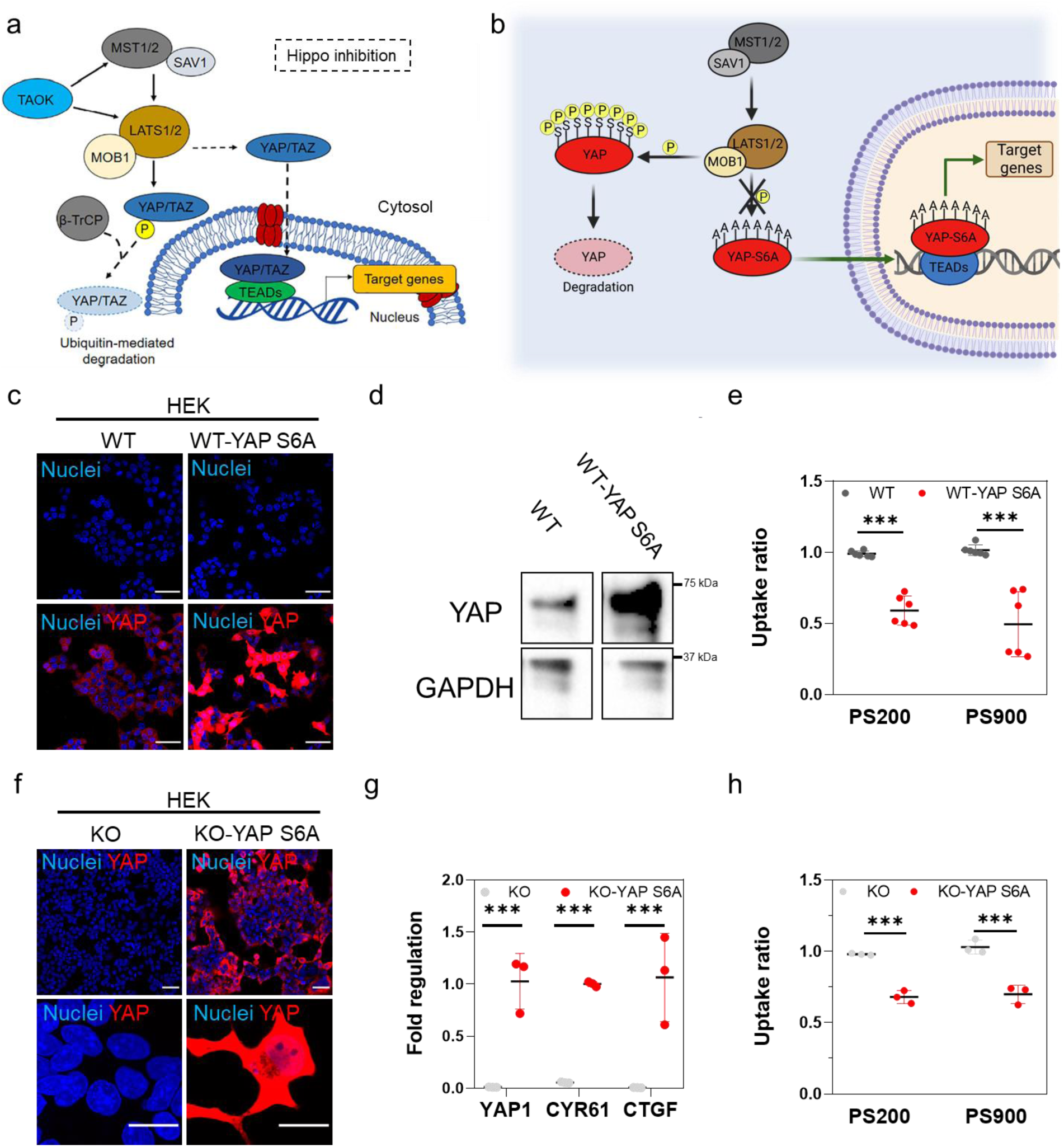
YAP overexpression or restoration in HEK 293T cells decreases their association with nanoparticles. (a) When the Hippo pathway is active, MST1/2 (STE20-like protein kinase 1/2) and SAV1 (protein salvador homolog 1) complex activates LATS1/2 (large tumor suppressor homolog 1/2) that phosphorylates, in association with MOB1 (MOB kinase activator 1), YAP, thus promoting its degradation. Conversely, when the Hippo signaling is inactive, YAP shuttles into the nucleus where it binds to TEADs (TEA domain transcription factor family members) and regulates the transcription of genes involved in cell proliferation, survival, and migration.^33^ TAOK, Serine/threonine-protein kinase TAO1; β-TrCP, Beta-transducin repeats-containing proteins; TAZ, Transcriptional coactivator with PDZ-binding motif. (b) Schematic representation of the constitutively active translocation of mutant YAP S6A to the cell nucleus. Owing to the substitutions of serine residues with alanine residues in six different positions (S61A, S109A, S127A, S128A, S131A, S136A, S164A, and S381A), YAP-S6A cannot be phosphorylated by upstream kinases, mainly belonging to the Hippo pathway (LATS1/2 kinases and scaffolding protein MOB1). Created with Biorender.com. (c) Representative confocal images of WT and WT HEK cells transfected with YAP S6A (WT-YAP S6A). Cells were decorated with DAPI (blue) and YAP (red). Scale bars: 50 μm. (d) Western blot analysis of the levels of YAP protein in WT HEK cells and WT-YAP S6A cells. Glyceraldehyde 3-phosphate dehydrogenase (GAPDH) was used for protein loading normalization. (e) Uptake ratios of PS200 and PS900 by WT HEK or WT-YAP S6A cells after incubation for 4 h. Statistical analysis was performed using two-way ANOVA followed by Sidak’s multiple comparison test; *n* = 6; \*\*\**p* < 0.001. (f) Confocal images of YAP −/− HEK (KO) cells and YAP −/− HEK cells transfected with YAP S6A (KO-YAP S6A). Scale bars: 50 and 10 µm in the lower and higher magnification images, respectively. (g) RT-qPCR analysis of YAP1, CYR61, and CTGF in KO and KO-YAP S6A cells. Statistical analysis was performed using multiple *t*-test; *n* = 3; \*\*\**p* < 0.001. (h) Uptake ratios of PS200 and PS900 in KO or KO-YAP S6A (red) cells. Statistical analysis was performed using two-way ANOVA followed by Sidak’s multiple comparison test; *n* = 3; \*\*\**p* < 0.001.

YAP expression in transfected WT HEK cells was first confirmed using confocal imaging (Fig. 4c; Fig. S6a) and western blot analysis (Fig. 4d). An empty vector was used as control. Subsequently, the transfected cells were incubated with PS200 or PS900 for 4 h. In line with the hypothesis that YAP activity impairs nanoparticle association, WT-YAP S6A HEK cells exhibited reduced nanoparticle uptake relative to the WT HEK control cells, as indicated by the uptake ratio obtained by flow cytometry (Fig. 4e; Fig. S6b,c). To further support these results, YAP −/− HEK cells were transfected with the plasmid carrying YAP S6A, and the expression of the protein was confirmed *via* confocal imaging (Fig. 4f). We note that the reintroduction of YAP in the KO cells increased the mRNA levels of CYR61 and CTGF, two of the main transcriptional targets of YAP, as assessed by RT-qPCR (Fig. 4g). Additionally, the reintroduction of YAP in YAP −/− HEK significantly decreased the uptake ratio of the nanoparticles by the cells (Fig. 4h), similar to what was observed with the overexpression of the protein in WT HEK cells.

Together, these results corroborate the key role of YAP in regulating the interactions between HEK cells and nanoparticles, suggesting that varying levels of protein activity may lead to different outcomes in this process. This opens the possibility of modulating YAP expression to enhance nanoparticle delivery.

### Pharmacological inhibition of MST1/2 kinase decreases cell–nanoparticle interactions in HEK 293T cells

Given that YAP-induced expression leads to a reduction in nanoparticle association, herein we sought to determine whether the pharmacological inhibition of the Hippo pathway kinase STE20-like protein kinase 1/2 (MST1/2) using the exogenous inhibitor XMU-MP1 could modulate cell interactions with nanoparticles.

XMU-MP1 is a small molecule inhibitor that blocks MST1/2 kinase, thereby preventing the activation of LATS1/2 and promoting YAP activation downstream of the Hippo pathway, leading to its nuclear shuttling (Fig. 5a).^37^ To assess the effect of XMU-MP1 treatment on YAP expression, WT HEK cells were treated with 6 μM of XMU-MP1 and the levels of the proteins downstream of the Hippo pathway were evaluated at different time points. Western blot analysis revealed a time-dependent decrease of MST1 (whereas the overall level of MST2 remained stable; Fig. S7) and a decrease of pospho-MOB1 (p-MOB1, Fig. 5b; Fig. S7) in cells treated with XMU-MP1. This result indicates that treatment with XMU-MP1 is effective in reducing the activity of the target kinase. Importantly, this treatment did not induce significant toxicity in WT HEK cells, as confirmed by a live/dead assay (Fig. S8).

**Fig. 5.**
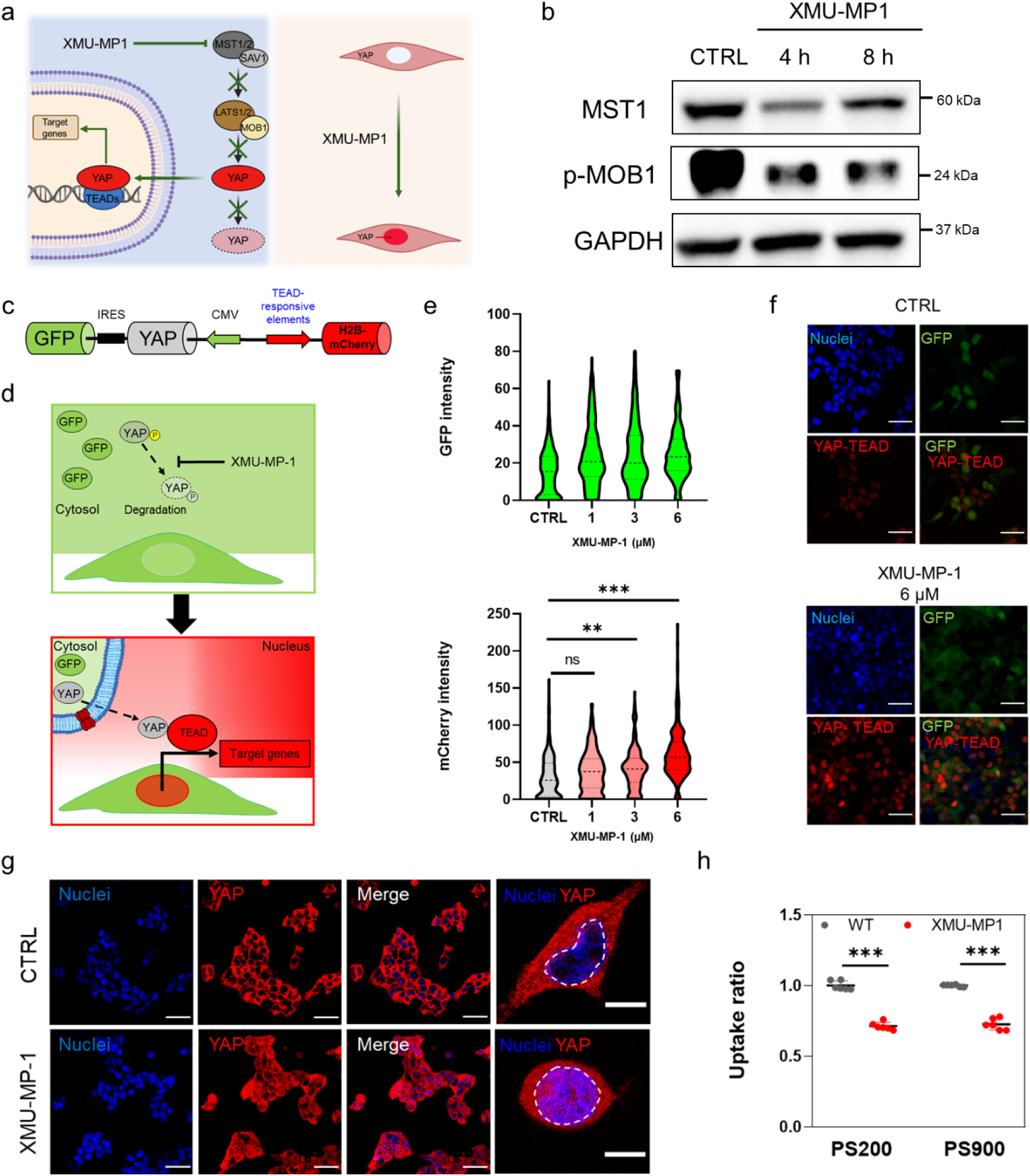
Suppression of the Hippo pathway with MST1/2 inhibitor XMU-MP1 in HEK 293T cells increases YAP activity and reduces cell–nanoparticle interactions. (a) Schematic representation of the effect of XMU-MP1 treatment. The drug inhibits the Hippo pathway by blocking the activity of upstream kinase MST1/2, thus suppressing the degradation of YAP and promoting its shuttling into the nucleus. Created with Biorender.com (b) Western blot showing the levels of MST1 and p-MOB1 in untreated HEK 293T cells (CTRL) or HEK 293T cells treated for 4 or 8 h with 6 µM XMU-MP1 inhibitor. GAPDH was used for protein loading normalization. (c) Scheme of the reporter construct used in this study, as described by Maruyama et al.^38^ FLAG-His6-YAP1 (FH-YAP1) gene is followed by IRESs and GFP gene are cloned under CMV promoter. Histone 2B-mCherry (H2B-mCherry) gene is regulated under the TEAD-responsive element. (d) Schematic showing that treatment with XMU-MP1 increases the H2B-mCherry signal, as the YAP-mediated TEAD transcriptional activity is promoted owing to inhibition of the activity of upstream kinase MST1/2 of the Hippo pathway. (e) (Top) Violin plot of the GFP signal intensity in HEK 293T cells treated with increasing concentrations of XMU-MP1 for 8 h (top). (Bottom) Violin plot of the signal intensity of H2B-mCherry in HEK 293T cells treated with increasing concentrations of XMU-MP1 for 8 h. Statistical analysis was performed using one-way ANOVA followed by Tukey’s multiple comparison test; *n* > 200 cells; ns, nonsignificant; \*\**p* < 0.01; \*\*\**p* < 0.001. (f) Representative confocal images of untreated WT HEK cells (CTRL) and WT HEK cells treated with 6 μM XMU-MP1 for 8 h. The green signal comes from GFP coexpressed with YAP, whereas the red signal comes from the YAP-TEAD-mediated gene transcription (mCherry). Cells are stained postfixation with DAPI (Merge, blue). Scale bars: 100 μm. (g) Representative confocal images of untreated HEK 293T cells (CTRL) and HEK 293T cells treated with 6 μM XMU-MP1 for 8 h. Cells are stained with DAPI (blue) and for YAP (Alexa Fluor 555, red). The magnified images show single untreated and treated cells with the nuclei delimited by the white dashed lines. Scale bars: 100 and 10 μm for the low and high magnification images, respectively. (h) Uptake ratios of PS200 and PS900 in untreated HEK 293T cells (CTRL) or HEK 293T cells treated with 6 μM XMU-MP1 for 8 h and incubated with the particles for 4 h (after 4 h of treatment with the inhibitor). Statistical analysis was performed using two-way ANOVA followed by Sidak’s multiple comparison test; *n* = 6; \*\*\**p* < 0.001.

Studies were then conducted to gain spatio-temporal insights into YAP activation following XMU-MP1 treatment and to evaluate the potential to finely modulate YAP activity, and consequently cell–nanoparticle interactions, using this inhibitor. WT HEK cells were transfected with a YAP transcriptional reporter constituted by mCherry-fused histone2B (H2B-mCherry) under the TEAD-responsive promoter and FLAG-tagged YAP1 linked to green fluorescent protein (GFP) *via* the internal ribosome entry site (IRES) under the cytomegalovirus promoter (CMV), as previously reported (Fig. 5c).^38^ Upon transfection with the reporter, YAP levels could be detected through the green signal resulting from GFP coexpression. In contrast, the transcriptional activity of YAP was identified by the red signal, which arises from YAP translocating to the nucleus and transcribing the YAP-TEAD-mediated reporter (as depicted in Fig. 5d).

After transfection with the reporter, WT HEK cells were sorted and incubated with XMU-MP1. A stable green signal was observed from the GFP in the transfected cells and treatment with the inhibitor for 8 h determined a concentration-dependent increase of mCherry signal (Fig. 5e). This result indicates that treatment with XMU-MP1 effectively activates YAP-TEAD transcriptional activity in the cells owing to its inhibitory activity on MST1/2 kinase (as demonstrated by western blot in Fig. 5b) and consequently its nuclear shuttling and transcriptional activity upon interaction with TEAD (Fig. 5f; Fig. S9). Treatment of HEK 293T cells at the highest concentration of XMU-MP1 studied (i.e., 6 μM) for 8 h resulted in the translocation of YAP into the nucleus, as confirmed by confocal analysis (Fig. 5g; Fig. S10a-d. Noteworthy, flow cytometry analysis showed a decrease in cell–nanoparticle association after incubation for 4 h with the inhibitor for both PS200 and PS900 (Fig. 5h; Fig. S10e,f). Despite the overall level of YAP remaining constant while the level of p-YAP varied (Fig. S11), our functional activity assay and the protein colocalization study demonstrated a significant increase in YAP activity upon XMU-MP1 treatment.

Collectively, the findings demonstrate a strong correlation between the modulation of YAP activity and the association of nanoparticles with cells. By tuning the functions of YAP, the Hippo pathway and the related mechanobiology pathways may represent a promising strategy to improve our understanding of the mechanisms involved in cell–nanoparticle interactions and enhance the design and the delivery efficiency of nanoparticles.

Noteworthy, the current findings were consistent in terms of cell–nanoparticle interactions with our previous findings of studies involving CAL51. However, some important differences were noted. The activity of YAP and the overall level of the protein in HEK 293T cells were lower than in CAL51 cells. The depletion of YAP did not result in a range of effects in HEK cells as those observed in CAL51 cells. Our study suggests the predominance of YAP activity in nanoparticle internalization over cell mechanics, as the physical properties of these cells, such as surface area, membrane stiffness, adhesion and assembly of focal adhesion, remain unchanged after YAP depletion in HEK cells.

## Conclusions

We demonstrated that depleting YAP in HEK 293T cells led to an increase in nanoparticle uptake, independently of nanoparticle size, thus underscoring the critical role of YAP activity in this process. This phenomenon highlights a potential mechanotargeting effect, where the cell mechanical response plays a prominent role in processing bio–nano interactions. Substrate properties and cell mechanics are often closely interconnected and have been shown to influence bio–nano interactions and nanoparticle uptake processes.^39^

Through RNA-seq analysis, we showed that YAP was involved in the transcription of genes related to membrane organization. WT HEK cells exhibited an extensive network of components at the cell membrane level, whereas YAP −/− HEK cells displayed a looser network, indicating a lower level of membrane interconnections. Building on these findings and our previous data,^29^ which also showed a less interconnected network of membrane protein in YAP-depleted CAL51 cells, we hypothesize that the level of membrane organization in cells, which is highly dependent on YAP activity, impacts cell–nanoparticle association, thus shedding light on the potential impact of mechanobiology in shaping the dynamics of bio–nano interactions. While our results indicate the transcriptional role of the protein as the primary factor responsible for this phenomenon, the significantly lower YAP level in HEK cells compared to CAL51, and its relatively higher cytoplasmic localization in the former, may suggest that other mechanisms, influenced by the protein at different levels and potentially dependent on the cytosolic pool of YAP, could be involved and warrant further investigation.

Although further research is needed to unravel the intricate interplay between mechanosensing and nanomaterials, particularly in assessing its preclinical and clinical significance, our study offers a fundamental mechanism to explain nanoparticle entry into cells. A deeper understanding of cell mechanobiology pathways involved in bio–nano interactions and the search for new targets and drugs to modulate their functions could pave the way for the development of next-generation nanotherapies, tailored to address the challenges of cell targeting and selective drug delivery in nanomedicine.

## Experimental Section

### Materials

The following materials were used in the present study: Opti-Link carboxylate-modified particles (83000520100290 and W090CA, Thermo Fisher Scientific); Dulbecco’s modified Eagle medium (DMEM, Merck); TrypLE Express (12604013, Thermo Fisher Scientific); Opti-Mem^TM^ medium (31985062, Thermo Fisher Scientific); 6-well plate (30006, SPL Life Sciences); penicillin/streptomycin (97063-708, VWR); 12-well plate (30012, SPL Life Sciences), 24-well plate (30024, SPL Life Sciences); µ-slide 8-well glass bottom dish (80807, Ibidi), tissue culture dish 40 mm (93040, iBiotech Ltd.); Pha-488 (A12379, Thermo Fisher Scientific); WGA-647 (W32466, Thermo Fisher Scientific); WGA-488 (W11261, Thermo Fisher Scientific); pLX304 (Addgene plasmid # 25890; http://n2t.net/addgene:25890; RRID:Addgene_25890); YAP1 (S6A) - V5 in pLX304 (Addgene plasmid # 42562; http://n2t.net/ addgene:42562; RRID:Addgene_42562); FuGENE^TM^ HD transfection reagent (E2311, Promega); lipofectamine 3000 (L3000001, Thermo Fisher Scientific); 10% mini-protean TGX precast protein gel (4561033, Bio-Rad); protease and phosphatase inhibitor cocktails (PPC1010, Merck); RIPA buffer (R0278, Merck); horseradish peroxidase (HRP)-conjugated anti-rabbit and HRP-conjugated anti-mouse (RABHRP1 and RABHRP2, Merck); MOWIOL 4-88 reagent (475904, Merck); High Pure RNA Isolation Kit (11828665001, Roche); LIVE/DEAD viability/cytotoxicity Kit (L3224, Thermo Fisher Scientific); TAMRA cadaverine (92001, Biotium); Float-A-Lyzer G2 dialysis device (300 kDa cutoff; G235036, Merck); *N*-hydroxysuccinimide (804518, Merck); inhibitor XMU-MP1 (S8334, Selleckchem); *N*-(3-dimethylaminopropyl)-*N*′-ethylcarbodiimide (EDC)-hydrochloride (59002, VWR); 4′,6-diamidine-2′-phenylindole dihydrochloride (10236276001, Merck); mouse anti-YAP (4912, Cell Signaling); mouse anti-Vinculin (V9131, Merck); rabbit anti-YAP (14074, Cell Signaling Technology; 1:1000); mouse anti-β-tubulin (T8328, Merck); rabbit anti-MST1 (3682, Cell Signaling Technology); rabbit anti-MST2 (3952, Cell Signaling Technology); rabbit anti-p-YAP (Ser397) (D1EZY, 13619, Cell Signaling Technology); rabbit anti-p-MOB1 (Thr35) (8699, Cell Signaling Technology); mouse anti-Glyceraldehyde 3-phosphate dehydrogenase (GAPDH)-peroxidase (69295, Merck); anti-rabbit immunoglobulin G (IgG), HRP-linked antibody (7074P2, Cell Signaling Technology); and Precision Plus Protein Dual Color Standards (1610374, Bio-Rad, 3 μL/well). All antibodies used for western blotting were diluted at a ratio of 1:1000 or 1:500 unless otherwise specified. Antibodies for immunohistochemistry were diluted at a ratio of 1:200.

### PS nanoparticle fluorescence labeling

Carboxylated PS nanoparticles with a diameter of 900 and 200 nm were labeled with TAMRA cadaverine or fluorescein cadaverine, as previously reported.^29^ Briefly, the particles were resuspended in 5 mL 2-(N-morpholino)ethanesulfonic acid (MES) buffer (50 mM, pH = 6.04) at a final concentration of 20 mg mL^−1^ and incubated with 52 μM *N*-(3-dimethylaminopropyl)-*N*′-ethylcarbodiimide hydrochloride and 5.2 μM *N*-hydroxysuccinimide for 1 h in an ice bath under magnetic stirring. TAMRA cadaverine or fluorescein cadaverine was then added, and the mixture was brought to a final concentration of 50 μM and left to react overnight at room temperature. Following reaction, the particles were collected, centrifuged (5000 g for 10 min for the 900 nm particles and 12,000 g for 15 min for the 200 nm particles), and washed five times with distilled water. Subsequently, the particles were dialyzed against distilled water for 72 h using a Float-A-Lyzer dialysis device (300 kDa cutoff).

### Generation of YAP mutant HEK 293T lines

The YAP −/− HEK 293T lines were generated using CRISPR/Cas9 technology as described previously.^28^ Briefly, a guiding RNA was designed to target exon 1 of the YAP1 gene, which is common in all nine YAP1 splicing variants. Two sets of complementary single-stranded DNA oligonucleotides (YAP1_R1: 50-CACCgtgcacgatctgatgcc-30, YAP1_R2: 50-AAACccgggcatcagatcgtgcac-30, YAP1_F1: 50-CACCGcatcagatcgtgcacgt-30, YAP1_F2: 50-AAACcggacgtgcacgatctgatgC-30) were then cloned into pSpCas9(BB)-2A-GFP (PX458) and transfected into HEK 293T cells using the FuGENE HD transfection reagent according to the manufacturer’s protocol. GFP-positive cells were sorted via fluorescence-activated cell sorting (MoFlo Astrios, Beckman Coulter, California, USA) as single cells and clonally propagated. Genomic DNA was sequenced from both sides to map the size of the deletion (sequencing primers: 50-gattggacccatcgtttgcg-30, 50-gtcaagggagttggagggaaa-30, 50-gaagaaggagtcgggcagctt-30, 50-gagtggacgactccagttcc-30).

### Cell culture

WT HEK 293T (gifted by Dr. V. Pekarik, Department of Physiology, Masaryk University, Brno, Czech Republic) and mutant cell lines were cultured in DMEM containing 10% inactivated fetal bovine serum, 1% penicillin-streptomycin, and 1% glutamine at 37 °C, 95% humidity, and 5% CO2. The cells were split every 2–3 days before reaching confluence, and only HEK 293T cells with a passage number of <10 were used for all experiments. Flow cytometry analysis was performed using unstained cells as a control. Cell viability was assessed using a LIVE/DEAD viability/cytotoxicity kit for mammalian cells, according to manufacturer’s instructions.

### AFM measurements

Cells were seeded onto a 40 mm tissue culture (TPP Techno Plastic Products, Trasadingen, Switzerland) at a concentration of 100,000 cells per dish for 24 h. Force maps were measured using a bio-AFM Nanowizard 4XP (Bruker-JPK, Germany) placed on a Leica DMi 8 inverted microscope with a 10× objective (Leica, Germany). A 5.73 µm melamine sphere (microParticles, Berlin, Germany) was attached to a soft tipless cantilever SD-qp-CONT-TL (NanoWorld, Neuchâtel, Switzerland) using epoxy resin. A petri dish containing either distilled water for calibration or cell culture was placed on a motorized stage preheated to 37 °C. Before each experiment, the laser reflection sum was maximized, and the laser detector was centered. Immediately after, probe sensitivity and stiffness were determined using the thermal noise method in Bruker-JPK software. Typical AFM settings were setpoint in the range of 0.2–0.8 nN relative to the baseline to maintain indentation depths up to 2.8 µm, *Z*-length at 10 µm, recording speed of 20 µm s^−1^, and sample rate at 5 kHz. Each force map consisted of 64×64 or 32×32 force–distance curves, covering an area of single or multiple cells, and Young’s modulus was calculated from the force–distance curves by fitting the DMT model^40^ in AtomicJ software.^41^

### Isolation of RNA and PCR analysis

Total RNA was isolated using a High Pure RNA Isolation Kit, according to the manufacturer’s protocol. Complementary DNA was synthesized using the RT^2^ First Strand Kit (SABiosciences Corporation, Frederick, USA). The expression levels of genes involved in ECM and cell adhesion were analyzed using RT^2^ Profiler PCR Arrays (Qiagen). RT-PCR was performed on the LigthCycler 480 Real-Time PCR System (Roche, Basel, Switzerland), and the cycling parameters were 1 cycle at 95 °C for 10 min, followed by 45 cycles at 95 °C for 15 s and 60 °C for 1 min. Normalization of gene expression levels was performed using an internal panel of housekeeping genes provided by the manufacturer. Genes with a high coefficient of variation among replicas or very low expression (35 < Ct < 40) were excluded from the analysis. The results are presented as heatmaps of quantification cycles (Ct) and graphs of mean ± standard deviation (s.d.) values of fold regulation, based on three samples per experimental condition.

### Cell transfection

Cells were transfected using lipofectamine 3000. The plasmids YAP1 (S6A)-V5 in pLX304 and pLX304 were obtained from Addgene as gifts from William Hahn (plasmid 42562) and David Root (plasmid 25890), respectively.^42^ HEK 293T cells were seeded onto a 6-well plate and transfected 24 h later with a preincubated mixture containing 250 μL Opti-MEM, 2.5 ng DNA, 7.5 μL lipofectamine 3000, and 5 μL P3000 reagent, added dropwise into each well. After 12 h, fresh medium was added, and cells were allowed to grow for another 8 h. The cells were then detached, seeded in a 24-well plate at a density of 200,000 cells/well, and cultured for an additional 12 h prior to incubation with nanoparticles.

### Generation of YAP reporter cell line

To produce second-generation lentiviral particles, HEK 293T cells were cotransfected with a three-plasmid combination: pMD2.G (Addgene #12259), psPAX2 (Addgene #12260), and pLL3.7 FLAG-YAP1-TEAD-P-H2B-mCherry (Addgene #128327) using FuGENE. The cell culture media was collected for 3 days, centrifuged and filtered to exclude cell debris, and pooled. Cells were incubated with media enriched in viral particles and supplemented with 10 μg mL^−1^ polybrene (Santa Cruz Biotechnology) for 6 h. The culture maintenance media was replaced daily and after 1 week, and cells were sorted for mCherry-positive cells on a MoFlo Astrios EQ (Beckman Coulter).

### Nanoparticle internalization studies

The day before the experiments, 200,000 cells were seeded in 500 μL of medium onto a 24-well plate. After 18 h, the cells were incubated with nanoparticles diluted at the desired concentration in the supplemented medium for 4 h (or according to the time indicated for each experiment), with 500 μL of nanoparticle suspension added per well. The samples were then processed for downstream flow cytometry and confocal analysis according to the following protocols: for cytofluorimetric analysis, WT or YAP −/− HEK 293T cells were cultured on 24-well plates and left for 12 h to adhere, then incubated with PS200 or PS900 for 4 h. For the experiments with the XMU-MP1, cells were treated with the desired concentration of inhibitor for 4 h, before the addition of PS200 or PS900 to the medium containing XMU-MP1 for additional 4 h. The medium was removed, 200 μL of TrypLE Express enzyme was added to each well for 5 min, and the cells were detached and washed three times until no particle signal was detected in the supernatant. The samples were analyzed using the FACSAria II flow cytometer (Becton Dickinson, USA), and plots were prepared with FlowJo software V10 (Tree Star, USA). For confocal laser scanning microscopy analysis, following incubation of cells with nanoparticles, the cells were detached, washed three times with phosphate-buffered saline (PBS), and left to adhere to a µ-slide 8-well glass bottom dish. After 4 h, the cells were processed according to the immunohistochemistry protocol described below.

### Immunohistochemistry and image analysis

For immunohistochemistry, 40,000 cells per well were seeded onto a µ-slide 8-well glass bottom dish for 24 h. After each relevant experiment, the medium was removed and the cells were washed with PBS. Before staining, the cells were fixed with 200 μL 4% paraformaldehyde in PBS for 15 min at room temperature, permeabilized with 0.1% Triton X-100 for 5 min, and then blocked with 2.5% bovine serum albumin (BSA) in PBS for 30 min. The primary antibodies were added in 200 μL PBS-BSA 2.5% solution and incubated for 2 h at room temperature or overnight at 4 °C. The secondary Alexa fluorochrome-conjugated antibodies were then added, and the cells were incubated in PBS. F-actin was stained with Pha-488 or Pha-647, the membrane was stained with WGA-488 or WGA-647, and the nuclei were counterstained with DAPI or Hoechst. The samples were embedded in Mowiol reagent and visualized with a Zeiss LSM 780 or Leica TCS SP8 X confocal microscope. *Z*-stacks were acquired with the optimal interval suggested by the software, and a maximum intensity algorithm was applied. Images were analyzed using ImageJ (http://rsb.info.nih.gov/ij/). The primary antibodies used were mouse anti-YAP (4912, Cell Signaling Technology) and mouse anti-Vinculin (V9131, Merck), diluted at a ratio of 1:200.

### Western blotting

Cells were lysed with RIPA buffer containing 1% protease and phosphatase inhibitor cocktails on ice, and then centrifuged at 14,000 *g* for 15 min at 4 °C. The supernatants were stored at 80 °C. Protein concentrations were determined using the bicinchoninic acid protein assay, and 20 μg of protein per sample was loaded onto 10% polyacrylamide gels and run at 100 V. Proteins were transferred to a polyvinylidene difluoride membrane using the Trans-Blot Turbo transfer system (Bio-Rad). The membranes were blocked with 5% BSA in tris-buffered saline - 0.1% Tween (TBST) and incubated with primary antibodies diluted in 5% BSA in TBST overnight at 4 °C. The membranes were then probed with HRP-linked secondary antibodies for 1 h at room temperature. Chemiluminescence was detected using the ChemiDoc imaging system (Bio-Rad), and band intensities were quantified using the Bio-Rad Image Lab software. The primary antibodies used were rabbit anti-YAP (1:1000), mouse anti-β-tubulin (1:1000), rabbit anti-MST1 (1:500), rabbit anti-MST2 (1:500), rabbit anti-pYAP S397 (1:500), rabbit anti-pMOB1 T35 (1:500), mouse anti-GAPDH-peroxidase (1:25,000); anti-rabbit IgG, HRP-linked antibody (1:1000).

### RNA-seq

For RNA-seq and data analysis, a library was prepared using the NEBNext Ultra II Directional RNA Library Prep Kit for Illumina with NEBNext Poly(A) mRNA Magnetic Isolation Module and NEBNext Multiplex Oligos for Illumina (Dual Index Primers Set 1). The kits were employed according to the manufacturer’s protocol, and 200–300 ng of total RNA was used as input to prepare the library. Sequencing was performed on an Illumina NextSeq 500 using the NextSeq 500/550 High Output v2 kit (75 cycles). Single-end 75bp sequencing was carried out in multiple sequencing runs until all samples had at least 30 million passing filter reads. Fastq files were generated using bcl2fastq software without any trimming. The quality of the raw sequencing data was assessed using FastQC (https://www.bioinformatics.babraham.ac.uk/projects/fastqc/) and aligned to the hg38 reference genome using TopHat2 aligner. Raw gene counts were calculated from reads mapping to exons, summarized by genes using the Ensembl 90 reference gene annotation (Homo sapiens GRCh38.p10, GTF) by HTSeq. Differential gene expression was determined using the DESeq2 Bioconductor package. Genes were considered differentially expressed if a Benjamini–Hochberg adjusted *P*-value was ≤0.05 and log2FC was ≥1. Biological term classification and gene cluster enrichment analysis were performed using the clusterProfiler package. All computations were performed using BioJupies.^43^ Enrichment analysis and ranking were performed using Enrichr, and the most significant annotations were downloaded from the available repository (https://maayanlab.cloud/Enrichr/).^44–46^ The GO categories were downloaded from the AmiGO 2 repository (https://amigo.geneontology.org).

### Image processing

Images were processed using FIJI, Imaris (version 10.0.1), and LAS X softwares. Using Imaris, stack images were first converted to imaris files with ImarisFileConverter, and 3D reconstruction was performed with the “volume rendering” function. Optical slices were obtained with the “orthoslicer” tool.

### Statistical analysis

Results were based on at least three replicates, and the data are presented as the mean ± s.d. The calculations were performed using GraphPad Prism v. 6.0 (San Diego, USA). For single-cell analysis, a minimum of 100 cells per sample were considered. The following statistical tests were used: unpaired *t*-test with Welch’s correction; multiple *t*-test; Kruskal–Wallis one-way ANOVA followed by Dunn’s multiple comparisons test; one-way ANOVA followed by Tukey’s multiple comparisons test; and two-way ANOVA followed by Sidak’s or Tukey’s multiple comparisons test. The appropriate statistical test was applied to the data as indicated in the figure captions for each experiment. A *P*-value of less 0.05 was considered statistically significant as denoted with asterisks (\**p* ≤ 0.05, \*\**p* ≤ 0.01, \*\*\**p* ≤ 0.001).

## Supporting information

Nanoscale Cassani et al., SI

## Author Contribution

**Conceptualization**: Marco Cassani, Giancarlo Forte

**Data curation**: Marco Cassani

**Formal Analysis**: Marco Cassani, Simon Klimovic, Jorge Oliver-De La Cruz, Jan Vrbsky

**Funding acquisition**: Marco Cassani, Giancarlo Forte, Frank Caruso

**Investigation**: Marco Cassani, Francesco Niro, Helena Durikova, Daniel Pereira-Sousa, Sofia Morazzo, Soraia Fernandes

**Methodology**: Jorge Oliver-De La Cruz, Jan Vrbsky, Simon Klimovic, Jan Pribyl, Tomas Loja

**Project administration**: Marco Cassani, Giancarlo Forte, Frank Caruso, Peter Skladal

**Supervision**: Marco Cassani, Giancarlo Forte, Frank Caruso

**Visualization**: Marco Cassani

**Writing – original draft**: Marco Cassani

**Writing -review & editing**: Marco Cassani, Soraia Fernandes, Giancarlo Forte, Jorge Oliver-De La Cruz, Frank Caruso

## Acknowledgments

We acknowledge František Foret for instrument access at the Department of Bioanalytical Instrumentation of the Institute of Analytical Chemistry of Brno, and Jana Bartoňová, Stefania Pagliari, and Vladimír Vinarský for scientific advice and technical assistance. We acknowledge the CF Genomics of CEITEC supported by the NCMG research infrastructure (LM2018132 funded by MEYS CR) Bioinformatics for their support with obtaining scientific data presented in this paper. We also acknowledge CIISB, Instruct-CZ Centre of Instruct-ERIC EU consortium, funded by MEYS CR infrastructure project LM2023042, and the European Regional Development Fund-Project “UP CIISB” (No. CZ.02.1.01/0.0/0.0/18_046/0015974) for financial support of the measurements at the CF Nanobiotechnology. We also thank Romana Vlčková, Hana Dulová, and Jana Vašíčková for their support on continuation of the study.

---

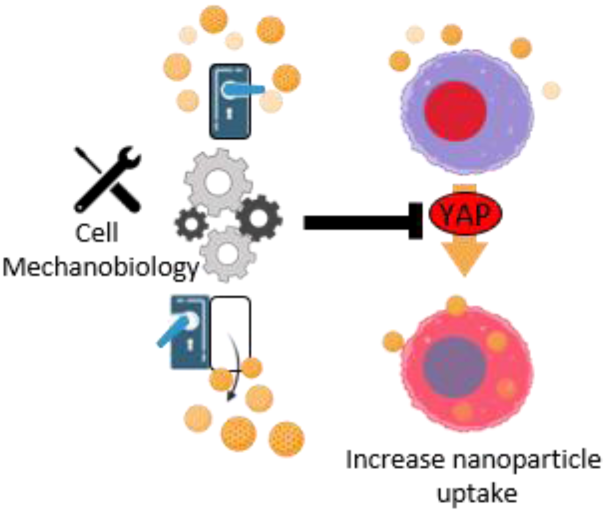
ToC. Tuning the mechanobiology of cells can be used to promote interactions between nanoparticles and the cell membrane. As a key factor in the regulation of cell mechanics, the inhibition of yes-associated protein (YAP) may be leveraged to optimize cell–nanoparticle interactions in promoting the delivery of nanomedicines.

## References

1. S. Wilhelm, A. J. Tavares, Q. Dai, S. Ohta, J. Audet, H. F. Dvorak and W. C. W. Chan, Nature Reviews Materials, 2016, 1, 16014.

2. R. van der Meel, E. Sulheim, Y. Shi, F. Kiessling, W. J. M. Mulder and T. Lammers, Nature Nanotechnology, 2019, 14, 1007–1017.

3. S. Fernandes, M. Cassani, S. Pagliari, P. Filipensky, F. Cavalieri and G. Forte, Current medicinal chemistry, 2020, 27, 7234–7255.

4. S. Fernandes, Advanced Science, 2023, DOI: 10.1002/advs.202305769.

5. S. E. Gratton, P. A. Ropp, P. D. Pohlhaus, J. C. Luft, V. J. Madden, M. E. Napier and J. M. DeSimone, Proceedings of the National Academy of Sciences of the United States of America, 2008, 105, 11613–11618.

6. J. P. Best, Y. Yan and F. Caruso, Advanced healthcare materials, 2012, 1, 35–47.

7. O. Shimoni, Y. Yan, Y. Wang and F. Caruso, ACS nano, 2013, 7, 522–530.

8. H. Sun, E. H. H. Wong, Y. Yan, J. Cui, Q. Dai, J. Guo, G. G. Qiao and F. Caruso, Chemical Science, 2015, 6, 3505–3514.

9. Y. Hui, X. Yi, D. Wibowo, G. Yang, A. P. J. Middelberg, H. Gao and C. X. Zhao, Science advances, 2020, 6, eaaz4316.

10. P. Guo, D. Liu, K. Subramanyam, B. Wang, J. Yang, J. Huang, D. T. Auguste and M. A. Moses, Nature Communications, 2018, 9, 130.

11. C. He, Y. Hu, L. Yin, C. Tang and C. Yin, Biomaterials, 2010, 31, 3657–3666.

12. Y. Jiang, S. Huo, T. Mizuhara, R. Das, Y. W. Lee, S. Hou, D. F. Moyano, B. Duncan, X. J. Liang and V. M. Rotello, ACS nano, 2015, 9, 9986–9993.

13. R. van der Meel, T. Lammers and W. E. Hennink, Expert opinion on drug delivery, 2017, 14, 1–5.

14. K. A. Dawson and Y. Yan, Nat Nanotechnol, 2021, 16, 229–242.

15. S. Sindhwani, A. M. Syed, J. Ngai, B. R. Kingston, L. Maiorino, J. Rothschild, P. MacMillan, Y. Zhang, N. U. Rajesh, T. Hoang, J. L. Y. Wu, S. Wilhelm, A. Zilman, S. Gadde, A. Sulaiman, B. Ouyang, Z. Lin, L. Wang, M. Egeblad and W. C. W. Chan, Nature Materials, 2020, 19, 566–575.

16. C. Huang, P. J. Butler, S. Tong, H. S. Muddana, G. Bao and S. Zhang, Nano letters, 2013, 13, 1611–1615.

17. D. Septiadi, F. Crippa, T. L. Moore, B. Rothen-Rutishauser and A. Petri-Fink, 2018, 30, 1704463.

18. Q. Wei, C. Huang, Y. Zhang, T. Zhao, P. Zhao, P. Butler and S. Zhang, Advanced materials (Deerfield Beach, Fla.), 2018, 30, e1707464.

19. D. Zhang, G. Wang, X. Yu, T. Wei, L. Farbiak, L. T. Johnson, A. M. Taylor, J. Xu, Y. Hong, H. Zhu and D. J. Siegwart, Nature Nanotechnology, 2022, 17, 777–787.

20. C. Sheridan, Nat Biotechnol, 2019, 37, 829–831.

21. S. Pagliari, V. Vinarsky, F. Martino, A. R. Perestrelo, J. Oliver De La Cruz, G. Caluori, J. Vrbsky, P. Mozetic, A. Pompeiano, A. Zancla, S. G. Ranjani, P. Skladal, D. Kytyr, Z. Zdráhal, G. Grassi, M. Sampaolesi, A. Rainer and G. Forte, Cell death and differentiation, 2021, 28, 1193–1207.

22. F. Martino, A. R. Perestrelo, V. Vinarský, S. Pagliari and G. Forte, Frontiers in physiology, 2018, 9, 824.

23. T. S. Gujral and M. W. Kirschner, Proceedings of the National Academy of Sciences of the United States of America, 2017, 114, E3729–e3738.

24. S. Dupont, L. Morsut, M. Aragona, E. Enzo, S. Giulitti, M. Cordenonsi, F. Zanconato, J. Le Digabel, M. Forcato, S. Bicciato, N. Elvassore and S. Piccolo, Nature, 2011, 474, 179–183.

25. T. Panciera, L. Azzolin, M. Cordenonsi and S. Piccolo, Nature reviews. Molecular cell biology, 2017, 18, 758–770.

26. Z. Pan, Y. Tian, C. Cao and G. Niu, Molecular cancer research : MCR, 2019, 17, 1777–1786.

27. F. Zanconato, M. Cordenonsi and S. Piccolo, Cancer cell, 2016, 29, 783–803.

28. G. Nardone, J. Oliver-De La Cruz, J. Vrbsky, C. Martini, J. Pribyl, P. Skládal, M. Pešl, G. Caluori, S. Pagliari, F. Martino, Z. Maceckova, M. Hajduch, A. Sanz-Garcia, N. M. Pugno, G. B. Stokin and G. Forte, Nature Communications, 2017, 8, 15321.

29. M. Cassani, S. Fernandes, J. Oliver-De La Cruz, H. Durikova, J. Vrbsky, M. Patočka, V. Hegrova, S. Klimovic, J. Pribyl, D. Debellis, P. Skladal, F. Cavalieri, F. Caruso and G. Forte, Advanced Science, n/a, 2302965.

30. P. Lajoie, L. D. Kojic, S. Nim, L. Li, J. W. Dennis and I. R. Nabi, Journal of cellular and molecular medicine, 2009, 13, 3218–3225.

31. H. De Belly, A. Stubb, A. Yanagida, C. Labouesse, P. H. Jones, E. K. Paluch and K. J. Chalut, Cell stem cell, 2021, 28, 273–284.e276.

32. M. Ruiz, R. Bodhicharla, E. Svensk, R. Devkota, K. Busayavalasa, H. Palmgren, M. Ståhlman, J. Boren and M. Pilon, eLife, 2018, 7.

33. M. Cassani, S. Fernandes, J. Vrbsky, E. Ergir, F. Cavalieri and G. Forte, Frontiers in bioengineering and biotechnology, 2020, 8, 323.

34. V. Rausch and C. G. Hansen, Trends in cell biology, 2020, 30, 32–48.

35. S. Verghese and K. Moberg, Frontiers in cell and developmental biology, 2019, 7, 384.

36. J. Rosenbluh, D. Nijhawan, A. G. Cox, X. Li, J. T. Neal, E. J. Schafer, T. I. Zack, X. Wang, A. Tsherniak, A. C. Schinzel, D. D. Shao, S. E. Schumacher, B. A. Weir, F. Vazquez, G. S. Cowley, D. E. Root, J. P. Mesirov, R. Beroukhim, C. J. Kuo, W. Goessling and W. C. Hahn, Cell, 2012, 151, 1457–1473.

37. F. Fan, Z. He, L. L. Kong, Q. Chen, Q. Yuan, S. Zhang, J. Ye, H. Liu, X. Sun, J. Geng, L. Yuan, L. Hong, C. Xiao, W. Zhang, X. Sun, Y. Li, P. Wang, L. Huang, X. Wu, Z. Ji, Q. Wu, N. S. Xia, N. S. Gray, L. Chen, C. H. Yun, X. Deng and D. Zhou, Science translational medicine, 2016, 8, 352ra108.

38. J. Maruyama, K. Inami, F. Michishita, X. Jiang, H. Iwasa, K. Nakagawa, M. Ishigami-Yuasa, H. Kagechika, N. Miyamura, J. Hirayama, H. Nishina, D. Nogawa, K. Yamamoto and Y. Hata, Molecular Cancer Research, 2018, 16, 197–211.

39. V. Panzetta, D. Guarnieri, A. Paciello, F. Della Sala, O. Muscetti, L. Raiola, P. Netti and S. Fusco, ACS biomaterials science & engineering, 2017, 3, 1586–1594.

40. B. V. Derjaguin, V. M. Muller and Y. P. Toporov, Journal of Colloid and Interface Science, 1975, 53, 314–326.

41. P. Hermanowicz, M. Sarna, K. Burda and H. Gabryś, 2014, 85, 063703.

42. X. Yang, J. S. Boehm, X. Yang, K. Salehi-Ashtiani, T. Hao, Y. Shen, R. Lubonja, S. R. Thomas, O. Alkan, T. Bhimdi, T. M. Green, C. M. Johannessen, S. J. Silver, C. Nguyen, R. R. Murray, H. Hieronymus, D. Balcha, C. Fan, C. Lin, L. Ghamsari, M. Vidal, W. C. Hahn, D. E. Hill and D. E. Root, Nature methods, 2011, 8, 659–661.

43. D. Torre, A. Lachmann and A. Ma’ayan, Cell systems, 2018, 7, 556–561.e553.

44. E. Y. Chen, C. M. Tan, Y. Kou, Q. Duan, Z. Wang, G. V. Meirelles, N. R. Clark and A. Ma’ayan, BMC Bioinformatics, 2013, 14, 128.

45. M. V. Kuleshov, M. R. Jones, A. D. Rouillard, N. F. Fernandez, Q. Duan, Z. Wang, S. Koplev, S. L. Jenkins, K. M. Jagodnik, A. Lachmann, M. G. McDermott, C. D. Monteiro, G. W. Gundersen and A. Ma’ayan, Nucleic acids research, 2016, 44, W90–97.

46. Z. Xie, A. Bailey, M. V. Kuleshov, D. J. B. Clarke, J. E. Evangelista, S. L. Jenkins, A. Lachmann, M. L. Wojciechowicz, E. Kropiwnicki, K. M. Jagodnik, M. Jeon and A. Ma’ayan, Current protocols, 2021, 1, e90.

