## Supplementary material for "Regulation of cell–nanoparticle interactions through mechanobiology": Nanoscale Cassani et al., SI

##### This PDF file includes:

- Figs. S1 to S11
- Reference S1

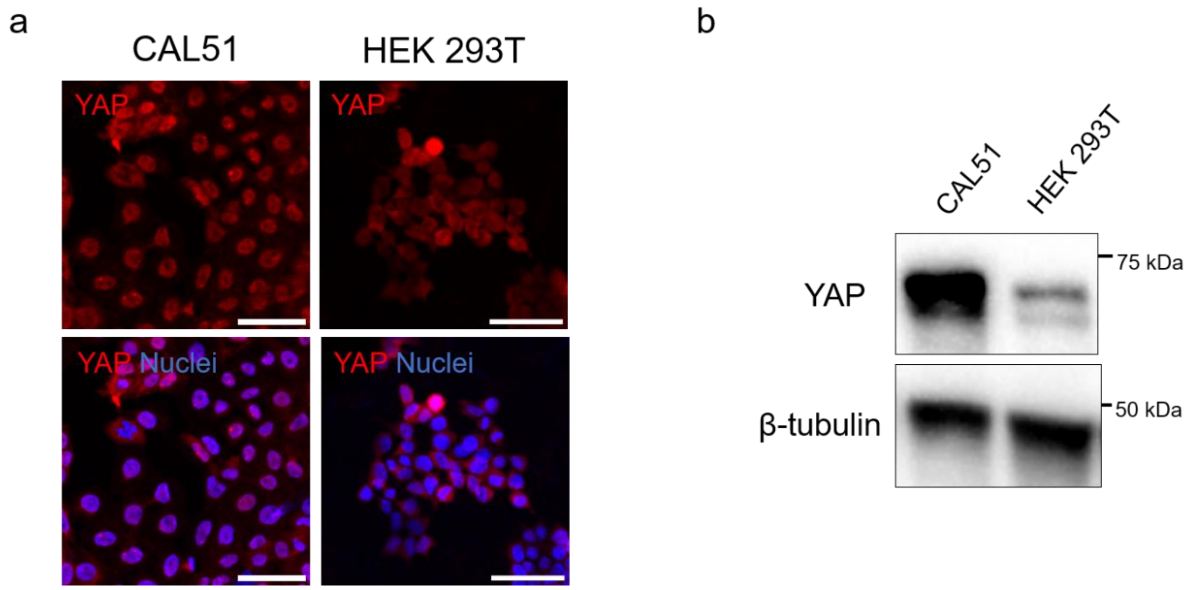


**Figure S1.** (a) Confocal images of CAL51 and HEK 293T cells stained for YAP (Alex Fluor 555, red) and DAPI (blue). Scale bars: 50 µm. (b) Western blot showing the levels of YAP in CAL51 and HEK 293T cells. YAP band in HEK 293T cells as shown in Figure 1a. β-Tubulin was used for protein loading normalization.

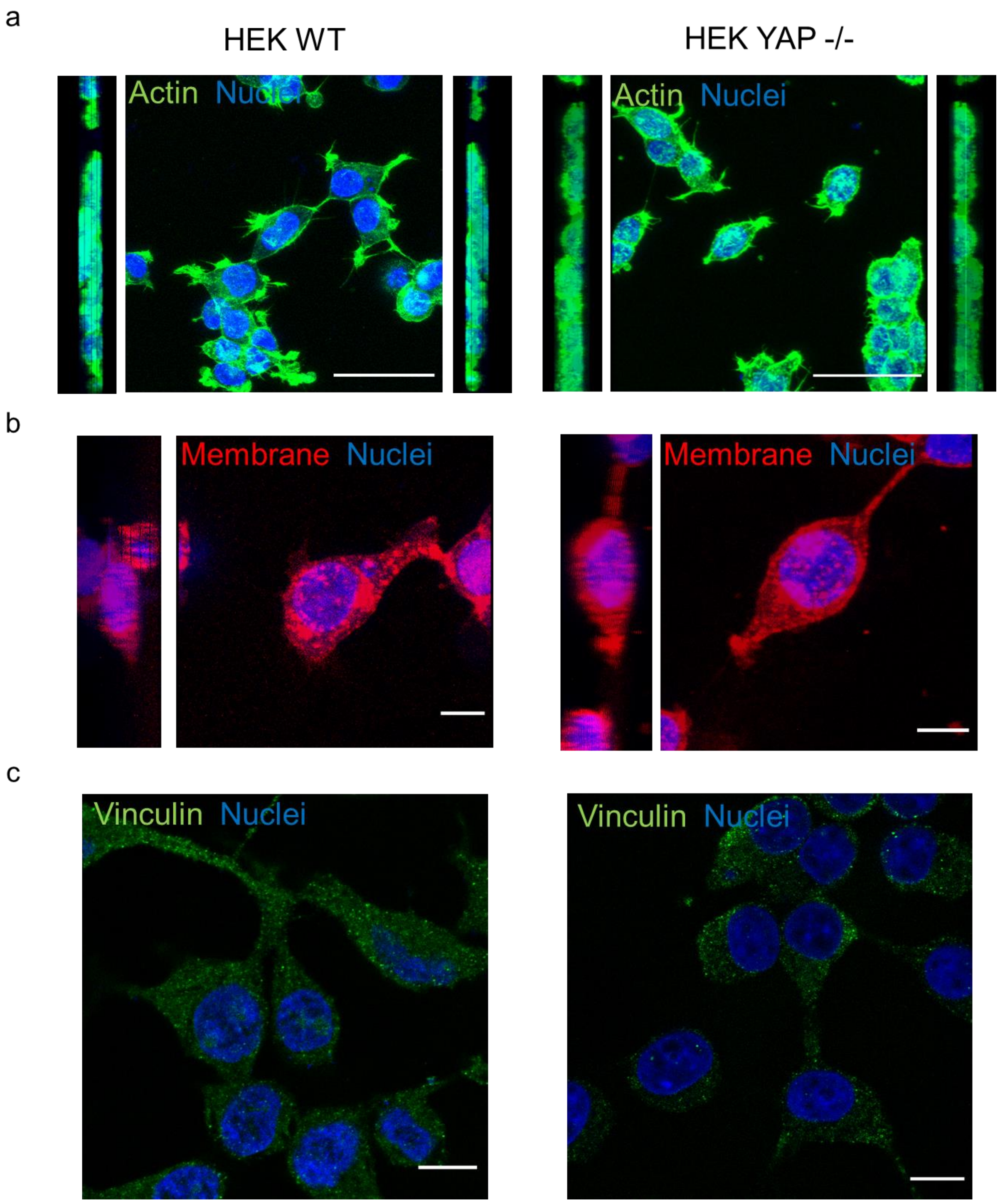


**Figure S2.** (a, b) 3D reconstruction of WT (a) and YAP −/− (b) HEK 293T cells. The lateral views are presented. The cells are stained with DAPI (blue), Pha-488 (green), WGA-647 (b, red) and vinculin (c, Alex Fluor 488). Scale bars: 50, 10, and 10 μm (a, b, and c, respectively).

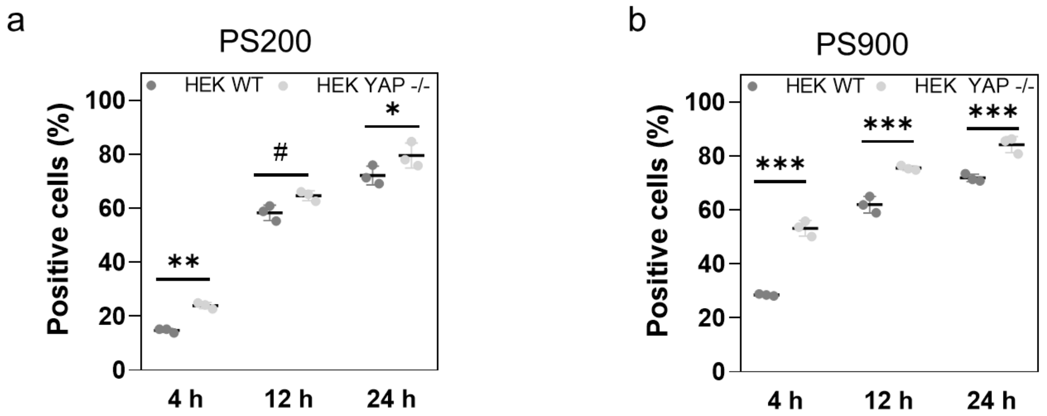


**Figure S3.** (a, b) Uptake of PS200 (a) and PS900 (b) in WT or YAP −/− HEK 293T cells following incubation for 4, 12, or 24 h. Statistical analysis was performed using two-way ANOVA followed by Tukey's multiple comparisons test; $n = 3$; ***$p < 0.001$; **$p < 0.01$; *$p < 0.05$; #$p = 0.0507$.

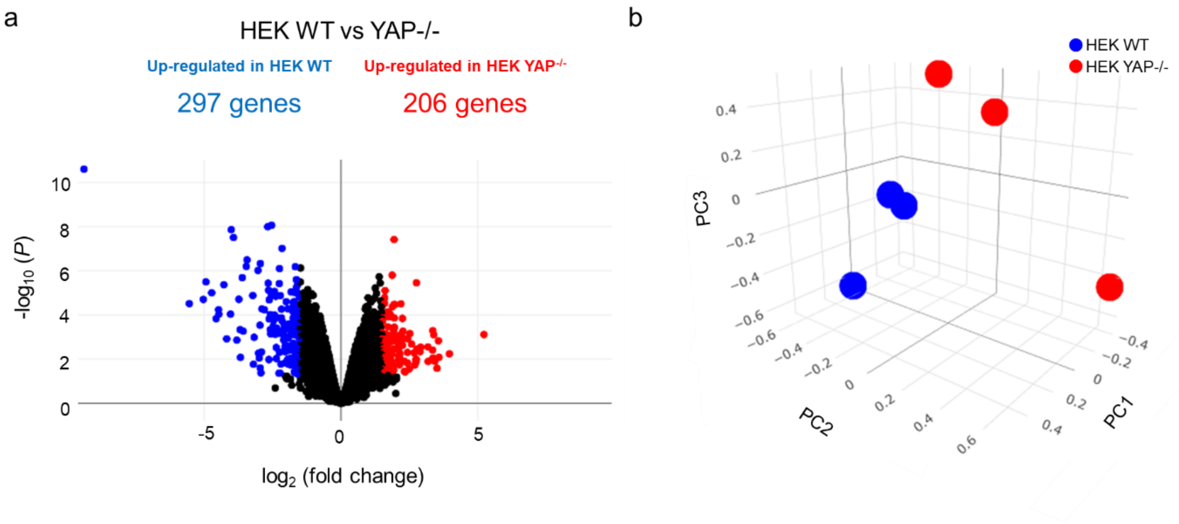


**Figure S4.** (a) Volcano plot showing differential gene expression in WT *vs* YAP −/− HEK 293T cells. Red markers indicate significantly upregulated genes and blue markers indicate downregulated genes; $N = 3$ (P adj < 0.05, log2FC > |1|). (b) 3D principal component (PC) analysis of RNA-seq in WT and YAP −/− HEK 293T cells. Red dots represent a sample of YAP −/− cells, whereas blue dots represent a sample of WT cells. $n = 4$ (P adj < 0.05, log2FC > |1|). The analysis was performed *via* Biojupies.[S1]

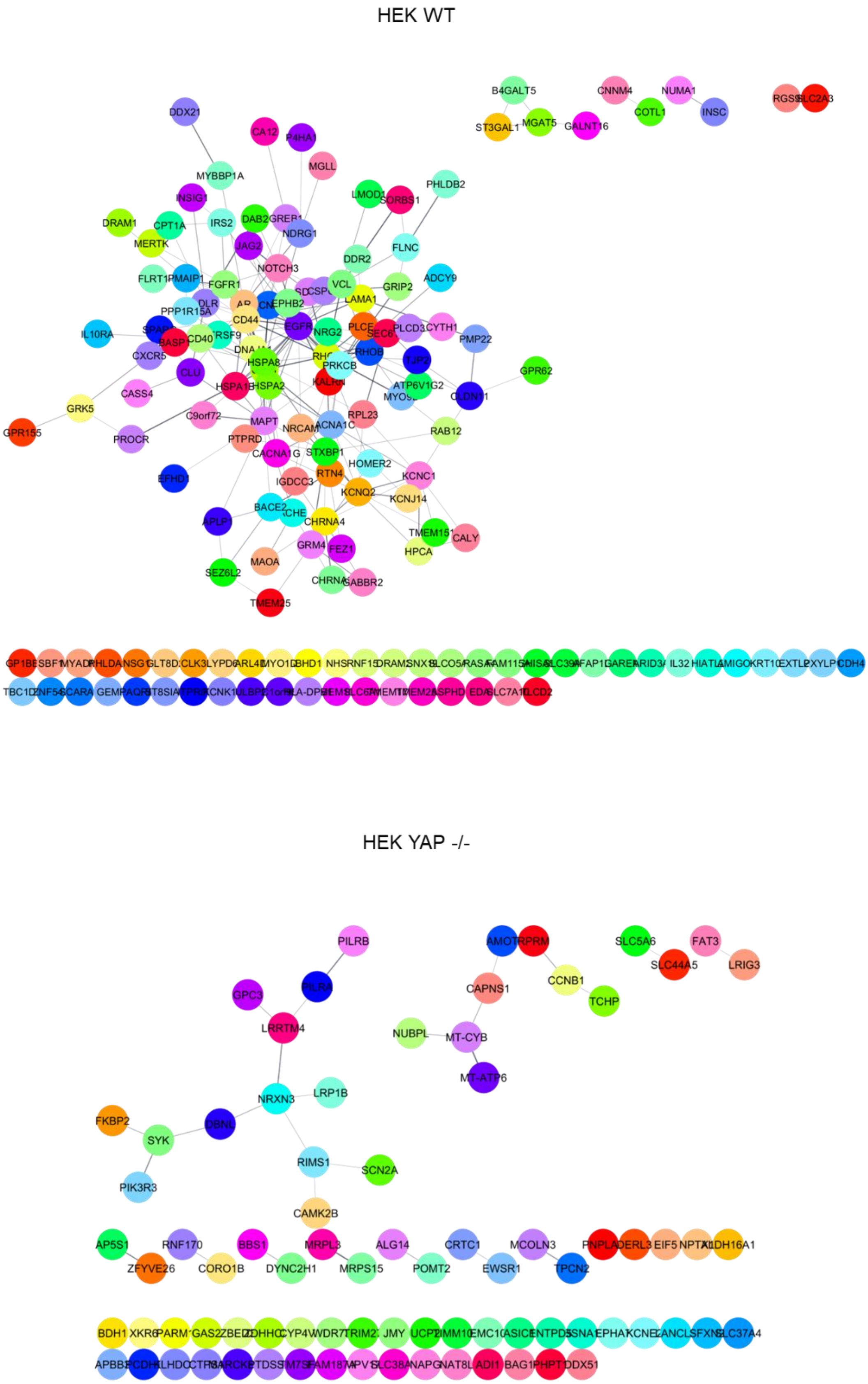


**Figure S5.** STRING PPI network of the differently expressed proteins involved in membrane organization (GO0016020) in WT and YAP −/− HEK cells obtained from Cytoscape (P adj < 0.05, log2FC > |1|, confidence cutoff 0.4).

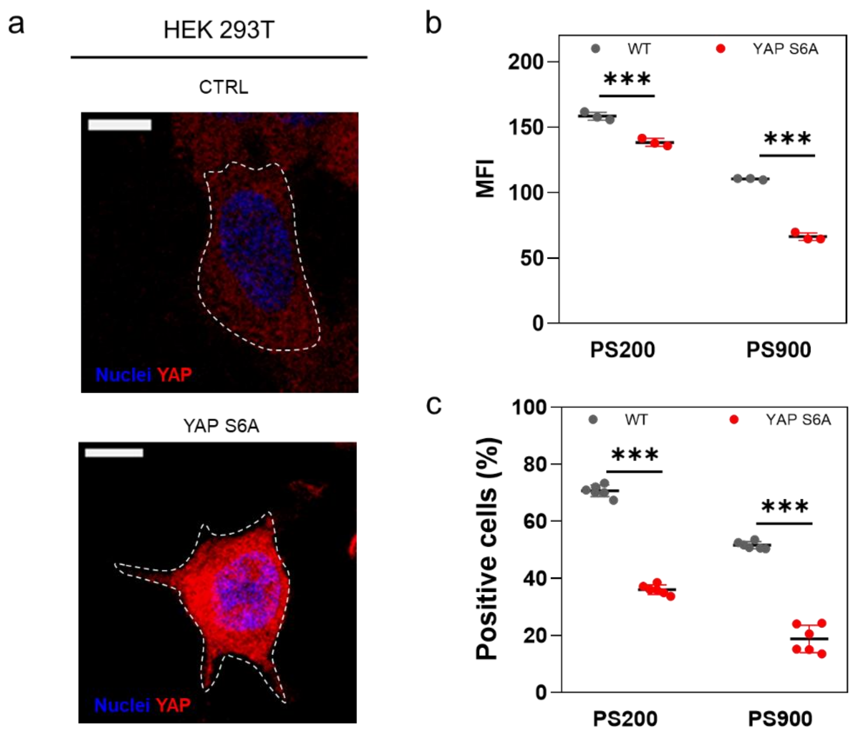


**Figure S6.** (a) Confocal images of WT HEK 293T cells (CTRL) and YAPS6A-transfected HEK 293T cells. Cells are stained for YAP (Alex Fluor 555, red) and DAPI (blue). Scale bars: 10 μm. (b) Median fluorescence intensity (MFI) of uptake of PS200 and PS900 in WT or YAPS6A cells following incubation for 4 h. Statistical analysis was performed using two-way ANOVA followed by Sidak's multiple comparisons test; $n = 6$; ***$p < 0.001$. (c) Uptake of PS200 and PS900 in HEK 293T WT or YAPS6A cells following incubation for 4 h. Statistical analysis was performed using two-way ANOVA followed by Sidak's multiple comparisons test; $n = 6$; ***$p < 0.001$.,

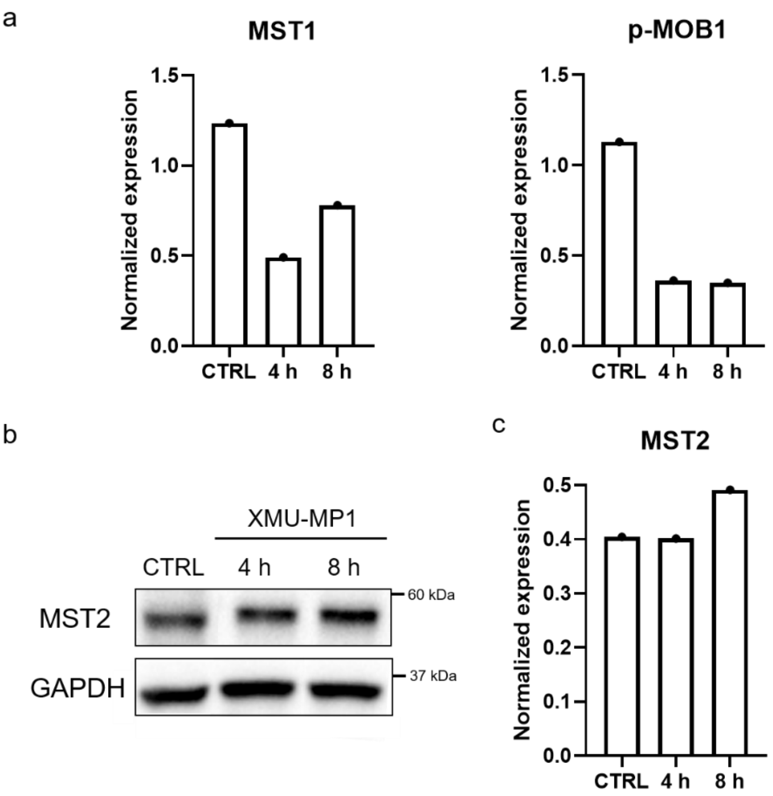


**Figure S7.** (a) Normalized expression of MST1 and p-MOB1 levels obtained from western blot analysis in untreated HEK 293T cells (CTRL) or HEK 293T cells treated for 4 or 8 h with 6 µM XMU-MP1 inhibitor. (b) Western blot showing the level of MST2 in untreated cells (CTRL) and HEK 293T treated for 4 or 8 h with 6 µM XMU-MP1 inhibitor. GAPDH was used for protein loading normalization. (c) Normalized expression of MST2 in untreated cells (CTRL) and HEK 293T cells treated for 4 or 8 h with 6 µM XMU-MP1 inhibitor.

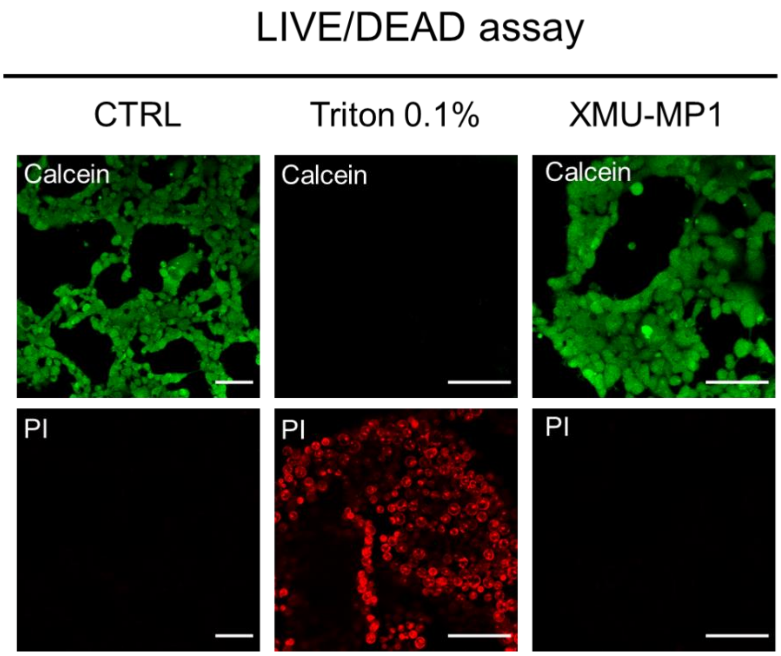


**Figure S8.** Live/dead assay performed on HEK 293T cells treated with XMU-MP1 for 8 h. As a control for cell death, cells were heated for 15 min with 0.1% Triton X-100 solution. Cells were excited with a 555 nm laser (propidium iodide, PI, red) and a 488 nm laser (calcein, green). Scale bars: 100 µm.

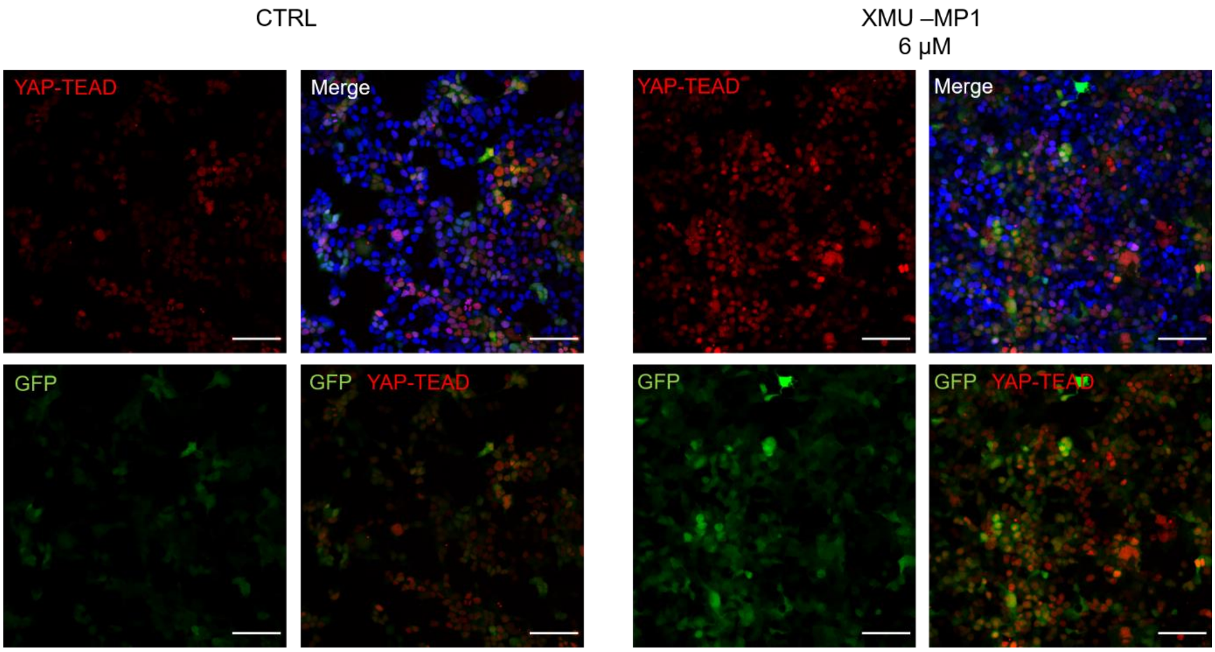


**Figure S9.** Representative confocal images of untreated WT HEK cells (CTRL) and HEK cells treated with 6 µM XMU-MP1 for 8 h. The green signal comes from the GFP protein coexpressed with YAP, whereas the red signal comes from the YAP-TEAD-mediated gene transcription (mCherry). Cells are stained postfixation with DAPI (Merge, blue). Scale bars: 100 µm.

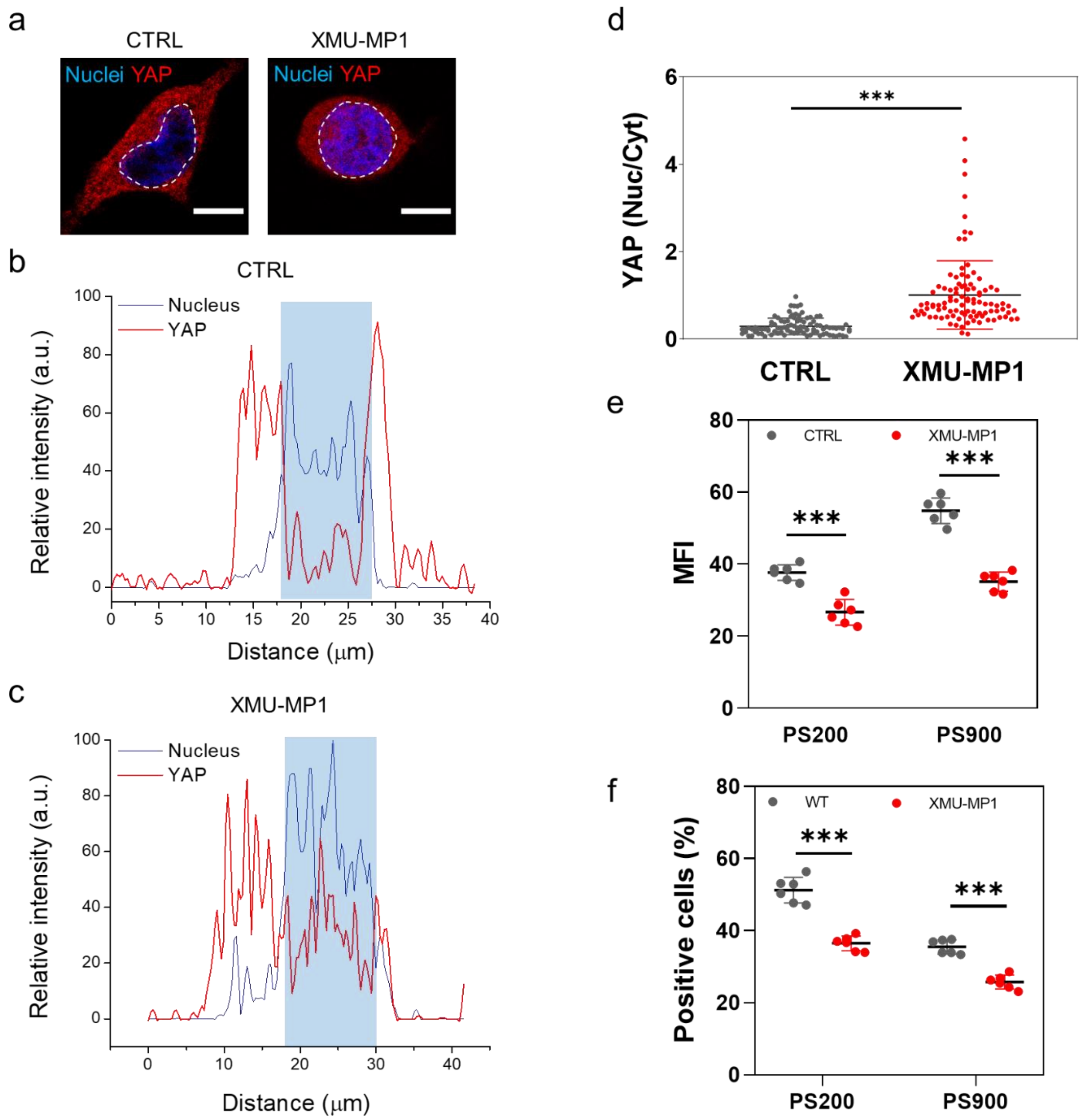


**Figure S10.** (a) Representative confocal images of untreated HEK 293T cells and HEK 293T cells treated with 6 μM XMU-MP1 for 8 h. The white dashed lines highlight the cell nuclei. Scale bars: 10 μm. The corresponding plot profiles of YAP signal intensity (Alex Fluor 555, red) colocalized with the nucleus (blue, DAPI) for the untreated cells (b) and XMU-MP1 treated cells (c) are also shown. (d) Dot plot representation of the YAP nuclear/cytoplasmic ratio in untreated HEK 293T cells and HEK 293T cells treated with 6 μM XMU-MP1 for 8 h. Statistical analysis was performed using unpaired *t*-test with Welch's correction; $n > 100$; ****p* < 0.001. (e) MFI of a 4-h uptake of PS200 and PS900 in untreated HEK 293T cells or HEK 293T cells treated with 6 μM XMU-MP1. Statistical analysis was performed using two-way ANOVA followed by Sidak's multiple comparisons test; $n = 6$; ****p* < 0.001. (f) Uptake of PS200 and PS900 in untreated HEK 293T cells or HEK 293T cells treated with 6 μM XMU-MP1 following incubation for 4 h. Statistical analysis was performed using two-way ANOVA followed by Sidak’s multiple comparisons test; $n = 6$; ***$p < 0.001$.

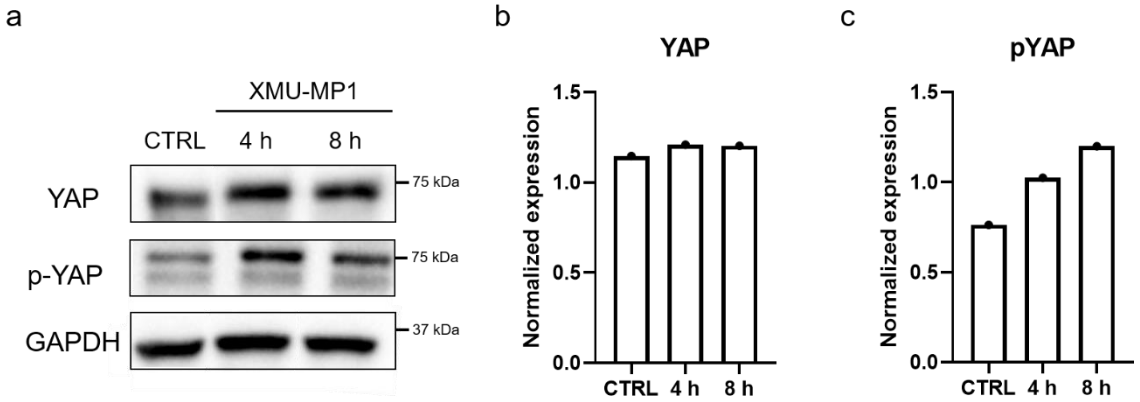


**Figure S11.** (a) Western blot showing the levels of YAP and p-YAP in untreated HEK 293T cells (CTRL) or HEK 293T cells treated for 4 or 8 h with 6 µM XMU-MP1 inhibitor. GAPDH was used for protein loading normalization. (b, c) Normalized expression of YAP (b) and p-YAP (c) in untreated HEK 293T cells (CTRL) and HEK 293T cells treated for 4 or 8 h with 6 µM XMU-MP1 inhibitor.

### References

S1. D. Torre, A. Lachmann and A. Ma'ayan, *Cell systems*, 2018, **7**, 556-561.e553.
